# Hedgehog signaling controls cytotoxic T cell migration in the tumour microenvironment

**DOI:** 10.1101/2024.08.14.607917

**Authors:** Chrysa Kapeni, Louise O’Brien, Dilyara Sabirova, Oliver Cast, Valentina Carbonaro, Stephen Clark-Leonard, Flavio Beke, Sarah McDonald, Kate Fife, Maike de la Roche

**Affiliations:** University of Cambridge, Cancer Research UK Cambridge Institute, Robinson Way, Cambridge CB2 0RE, UK; Histopathology Department, Cambridge University Hospitals NHS Foundation Trust, Hills Road, Cambridge CB2 0QQ, UK; Oncology Department, Cambridge University Hospitals NHS Foundation Trust, Hills Road, Cambridge CB2 0QQ, UK

## Abstract

Cytotoxic T lymphocytes effectively eliminate cancer cells. Their abundance in the tumour microenvironment is one of the strongest pan-cancer predictors of clinical response. Here, we show that Hedgehog (Hh) signaling regulates T cell migration into tumours.

Using conditional knockout mouse models of central Hh signaling components *Ihh*, *Smo* and *Gli1* in CD8 T cells, we show that *Smo* deletion greatly impairs the anti-tumour response *in vivo* due to diminished CD8 T cell migration into the tumour microenvironment. The migration defect is mediated exclusively by Smo, both in *in vivo* cancer models and *in vitro* migration assays. This effect is independent of the canonical Hh pathway and relies on the GPCR function of Smo to regulate the migration of murine and human CD8 T cells via RhoA.

Hh signaling is critical during embryonic development and adult stem cell homeostasis, but is also amplified in multiple cancer types. Hh inhibitors targeting SMO have been clinically-approved and shown efficacy in the treatment of Hh-driven basal cell carcinoma and medulloblastoma but have failed in clinical trials in other solid cancers with upregulated Hh signaling. We demonstrate that SMO inhibitors specifically decrease CD8 T cell migration into the tumour microenvironment, both in murine cancer models and resected BCCs from patients treated with the SMO inhibitor vismodegib, providing the first mechanistic explanation as to why Hh inhibitors have failed in solid cancers.

Our data establishes a novel link between Hh inhibition *in vivo* and the anti-tumour immune response and reveals a fundamental mechanism controlling T cell migration. The work provides the basis for improved Hh targeting approaches in the clinic and new entry points into enhancing migration in T cell therapies.

## INTRODUCTION

The Hedgehog (Hh) pathway orchestrates cell fate choices throughout metazoan development and adult tissue homeostasis by regulating tissue patterning, cell proliferation and differentiation. In canonical signaling, extracellular Sonic, Desert or Indian Hh ligands (Shh, Dhh, Ihh) bind to the transmembrane receptor Patched (Ptch) on Hh-responsive cells. Upon binding, Ptch releases its inhibition of the transmembrane protein Smoothened (Smo). Smo translocates into the primary cilium, where it activates the glioma-associated oncogene (Gli) transcription factors, Gli1, Gli2 and Gli3. These move into the nucleus and initiate a Hh-specific transcriptional programme^20^.

The pathway is also amplified in diverse human cancer types. Mutations in the pathway are oncogenic drivers in BCC^1, 2, 3, 4^, medulloblastoma^5, 6, 7, 8, 9, 10^ and a subset of rhabdomyosarcoma^11, 12^. In addition, the pathway is aberrantly upregulated in many human cancers including gastrointestinal cancers such as pancreatic and colorectal adenocarcinoma as well as lung, breast, prostate, and haematological malignancies (reviewed in^13^). As a result, Hedgehog inhibitors have emerged as promising anti-cancer therapeutics and clinically approved small molecule Hedgehog inhibitors have been the subject of multiple clinical trials (reviewed in^14^). Despite promising preclinical data, Hedgehog inhibitors have largely failed to meet primary endpoints in clinical trials of patients with colorectal^15^, pancreatic^16, 17, 18^ and prostatic adenocarcinomas^19^. Elucidating the mechanisms for this treatment failure is a high priority.

SMO inhibitors such as vismodegib and sonidegib (Erivedge and Odomzo, respectively) have been approved by the FDA and EMA for the treatment of non-resectable advanced (both drugs) and metastatic (only vismodegib) BCCs. Little is known about the effect of Hh inhibitors on the cells of the immune system. Previous work has shown that the pathway is required for CD8 cytotoxic T lymphocyte (CTL) killing^21^. In T cells, the pathway is activated upon engagement of the T cell receptor (TCR) and controls actin clearance and centrosome polarization, a process essential for the formation of the immunological synapse and the targeted release of cytotoxic granules^21^. This work suggested that a reason for the lack of efficacy of Hedgehog inhibitors in the clinic for many indications might be due to the inhibition of CTL killing, resulting in a diminished anti-tumour immune response. However, to date, no functional assessment of the anti-tumour T cell response during Hedgehog inhibition *in vivo* in the tumour microenvironment has been performed. In this study, we show that both pharmacological and genetic inhibition of Hh signaling *in vitro* and *in vivo* greatly impairs CTL migration into the tumour. Mechanistically, we show that the reduced CTL migration is not mediated by canonical Hh signaling and is therefore independent of the ligand Ihh and the transcription factor, Gli1, which is implicated in the formation of the immunological synapse. Instead, CTL migration in mouse and human is regulated non-canonically via the Hh signal transducer Smo and its GPCR function, which controls downstream levels of active RhoA. Finally, we are able to show that migration of cytotoxic T cells into BCCs is diminished when patients are treated with vismodegib.

## RESULTS

### Hh inhibitors in clinical trials for treatment of cancers with amplified Hh signaling have only benefited a subset of patients

To broadly assess the performance of Hh inhibitors in the clinic, we compiled the results of published clinical trials using these agents in cancer treatment. We compiled trials that used the SMO antagonists vismodegib^22, 23^, sonidegib^24^, and itraconazole^25, 26^. We also included the GLI inhibitor arsenic trioxide (ATO)^27^ **(Fig. 1, Suppl. Table 1)**. Tumours with a Hh signaling driver mutation, such as basal cell carcinoma (BCC) and SHH medulloblastoma, benefited the most from Hh inhibition and showed remarkable response rates ranging between 8-71%. Of note, when vismodegib was combined with the PD-1 antagonist Pembrolizumab in locally advanced BCC (laBCC), the treatment benefit was negated. In paediatric medulloblastoma, lung cancer, prostate cancer and acute myeloid leukaemia, Hh inhibitors failed to meet the primary endpoint. Critically, Hh inhibitor treatment was detrimental in patients with pancreatic, stomach and colorectal cancer compared to standard treatment alone **(Fig. 1, Suppl. Fig. 1)**.

**Figure 1:**
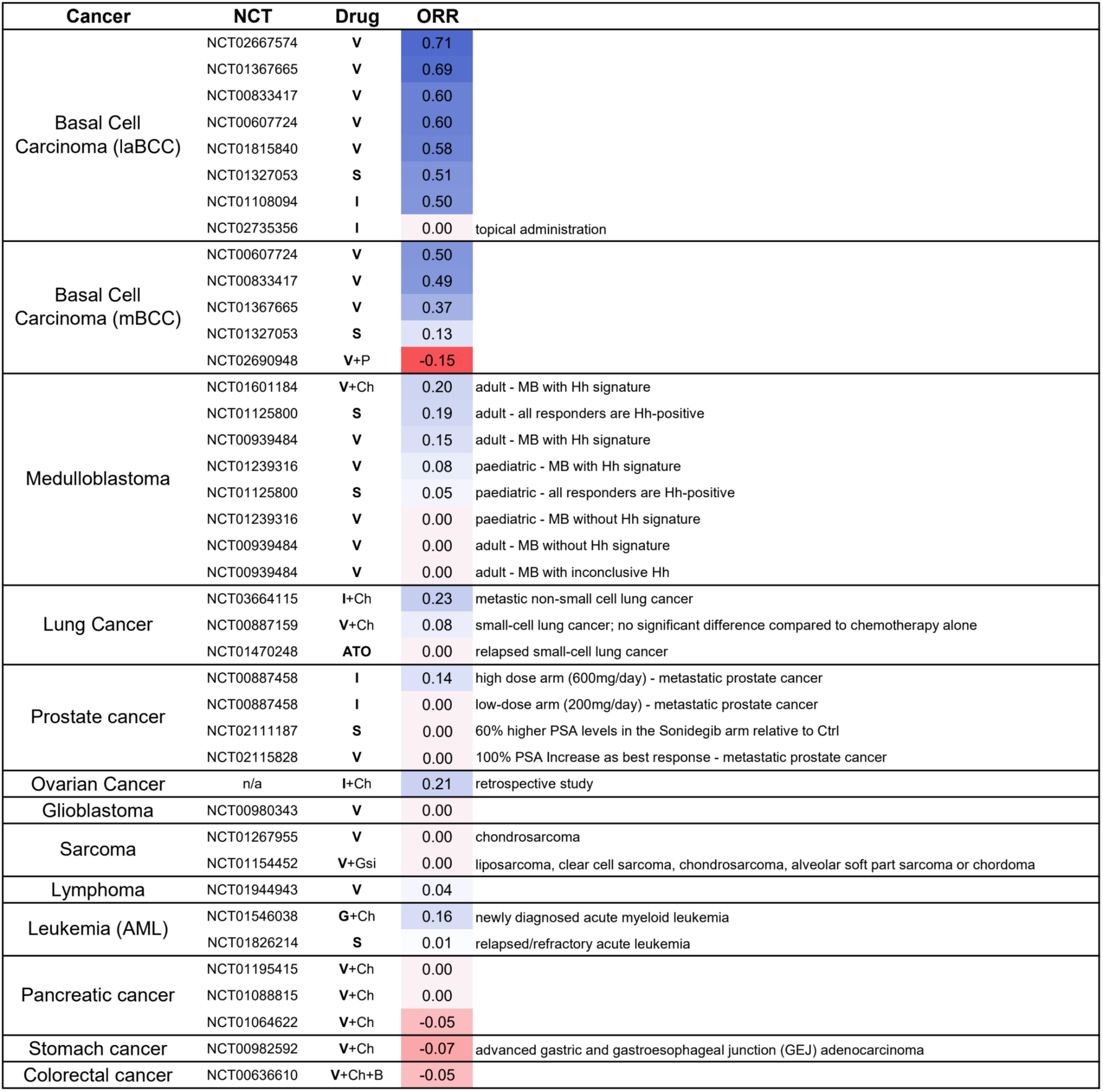
Clinical trials of Hedgehog inhibitors in cancer patients have shown low efficacy in non-Hh-driven malignancies. Clinical trials using Hh inhibitors in cancers with driver mutations in the Hh pathway (laBCC, mBCC, a subset of medulloblastoma patients) and cancers with an amplified Hh signature (lung, prostate, sarcoma, leukemia, pancreatic, stomach and colorectal). Overall Response Rate (ORR) shows the percentage of complete responders or partial responders. When indicated, Hh inhibitors were administered in addition to the current gold standard of treatment. In those cases, the ORR represents the difference between the response rate of the standard treatment alone versus standard treatment in combination with Hh inhibitor. Abbreviations: BCC: Basal Cell Carcinoma, laBCC: locally advanced BCC, mBCC: metastatic BCC, NCT: ClinicalTrials.gov identifier, V: vismodegib, S: sonidegib, I: itraconazole, ATO: arsenic trioxide P: pembrolizumab, Ch: chemotherapeutic agents (respective standard of care), Gsi: Gamma-secretase Inhibitor RO4929097, G: glasdegib, B: bevacizumab.

### Tumour growth in a mouse model of colorectal cancer is exacerbated upon sonidegib treatment

To investigate the lack of efficacy of Hh inhibitors in gastrointestinal cancers, we used a mouse model of colorectal adenocarcinoma. For this, we injected C57BL/6J mice subcutaneously with MC38 tumour cells and let the tumour establish for twelve days. Mice were then stratified into two equal groups and treated daily by oral gavage with Smo inhibitor sonidegib or carrier control up to day 24 **(Fig. 2A, B)**. Successful systemic inhibition of the Hedgehog pathway *in vivo* was confirmed by expression of the key Hedgehog target genes *Gli1* and *Ptch1*, which were both significantly reduced **(Fig. 2C)**. Strikingly, mice treated with sonidegib exhibited a doubling of tumour load as measured by size **(Fig. 2D)** and weight **(Fig. 2E)** and increased spleen weight.

**Figure 2.**
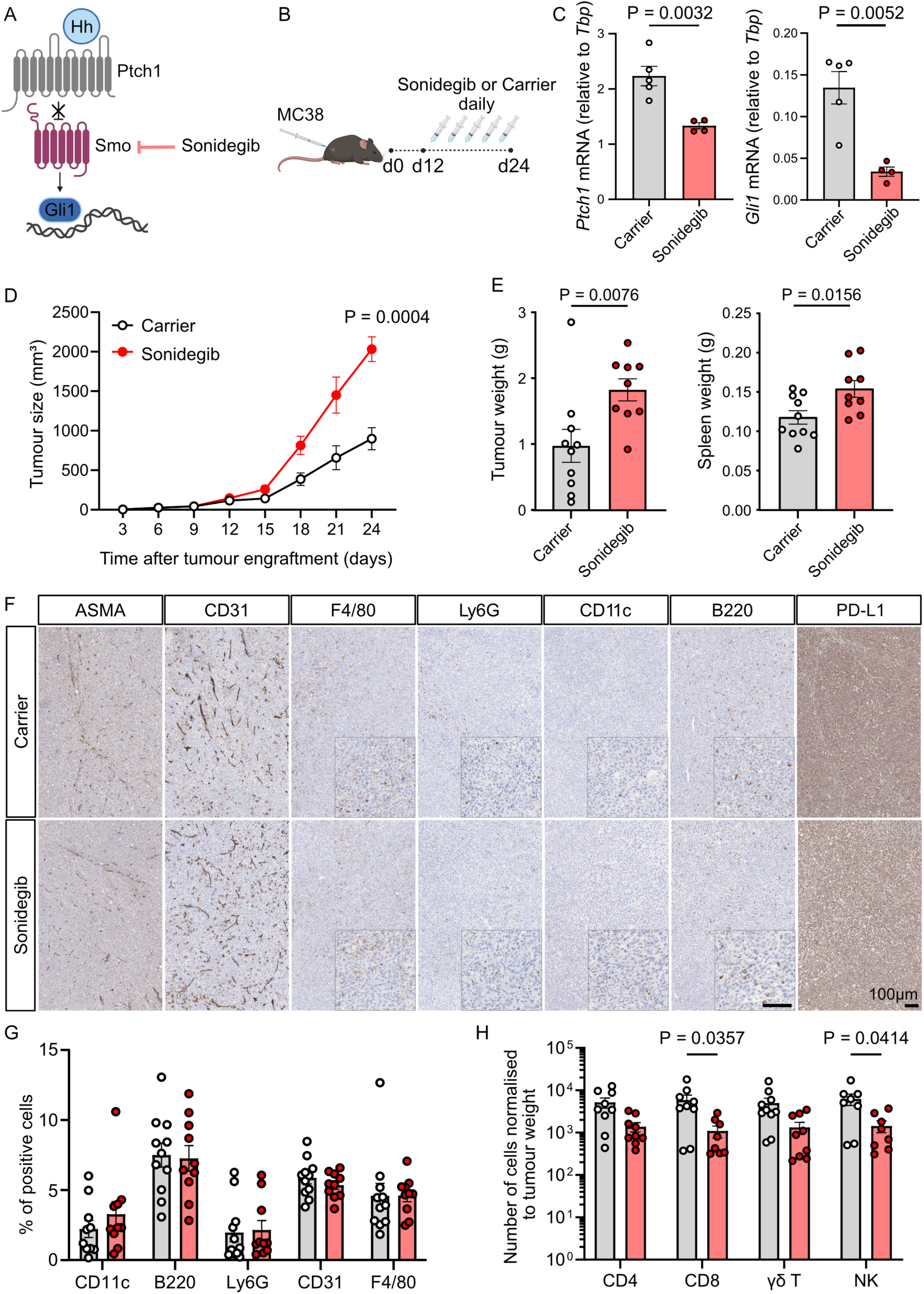
Treatment with clinically approved Hh inhibitor sonidegib exacerbates colorectal cancer growth. **(A)** Schematic of the canonical Hh signaling pathway. In the presence of Hh ligands, Ptch1 releases its inhibition of Smoothened (Smo), which leads to the activation of Gli transcription factors inducing a Hh-specific target gene programme. Sonidegib is an FDA/EMA-approved Smo-specific antagonist. **(B)** Experimental Design. C57BL/6J wildtype mice were subcutaneously injected with 0.5 × 10^6^ MC38 colorectal tumour cells on day(d)0. On d12, mice were stratified into two equal groups according to tumour size. Between d12 and d24, mice were treated daily with 20mg/kg sonidegib or carrier control by oral gavage. **(C)** mRNA levels of *Ptch1* and *Gli1* in the small intestine. One independent experiment, n=5 for carrier-treated mice, n=4 for sonidegib-treated mice, unpaired t-test, mean ± SEM. **(D)** Tumour growth was determined from caliper measurements. Two independent experiments, n=10 for carrier-treated mice, n=9 for sonidegib-treated mice, ordinary two-way ANOVA with Geisser-Greenhouse correction, mean ± SEM. **(E)** Tumour and spleen weight at endpoint (d24). Two independent experiments, n=10 for carrier-treated mice, n=9 for sonidegib-treated mice, unpaired Mann-Whitney test, mean ± SEM. **(F)** Representative images of MC38 tumours from mice treated with carrier (top row) or sonidegib (bottom row) and stained with antibodies against cell subsets of the tumour microenvironment as indicated. **(G)** Quantification of cell subsets in the tumour microenvironment by IHC staining pictures shown in (**F**). Two independent experiments, n=11 for carrier-treated mice, n=10 for sonidegib-treated mice, two-way ANOVA, mean ± SEM. **(H)** Numbers of CD4, CD8, gammadelta T cells and NK cells in tumour microenvironment assessed by flow cytometry and normalised to tumour weight. Two independent experiments, n=10 for carrier-treated mice, n=9 for sonidegib-treated mice, two-way ANOVA, mean ± SEM.

### Sonidegib treatment does not lead to drastic changes in tumour composition

To assess whether sonidegib treatment led to global changes in the tumour microenvironment, we stained for fibroblasts (ASMA), endothelial cells (CD31), macrophages (F4/80), myeloid cells (Ly6G), dendritic cells (CD11b) and B cells (B220) in tumour tissue sections by IHC **(Fig. 2F)**. Upon quantification, no significant differences were detected between sonidegib-treated versus carrier-treated tumour groups **(Fig. 2G)**. We also assessed the tumour cell composition by analysing bulk RNASeq data using ConsensusTME^28^ **(Supp. Fig. 2A)**. The analysis confirmed no significant changes in the tumour stroma, blood vessels or in the myeloid cell, B cell, and dendritic cell compartments.

### Cytotoxic T cell infiltration into the tumour is diminished upon sonidegib treatment

Despite no observational change in the overall tumour composition, a significant reduction in the infiltration of predominantly cytotoxic lymphocytes into the tumour mass was detected by flow cytometry **(Fig. 2H)** and bulk RNASeq (Consensus TME) **(Suppl. Fig. 2A)**. Cytotoxic CD8 T cells and NK cells in the TME of sonidegib-treated mice were 81% and 77% reduced, respectively. The levels of CD4 T cells and gammadelta T cells were also moderately reduced.

To investigate this further, we performed IHC staining for CD8 cells on tumour sections **(Fig. 3A, B)**. To assess the extent of CD8 T cell infiltration into the tumour mass, we used the imaging software HALO to segment the tumour in 100micron-wide concentric zones **(Suppl. Fig. 3A)**. Our analysis showed that not only were there more CD8 T cells (CD3+, CD8+) in carrier-treated tumours, but the CD8 T cells were also able to travel further towards the tumour centre compared to sonidegib-treated tumours **(Fig. 3C)**. This was also true for total CD3 T cells **(Suppl. Fig. 3B, C)**.

**Figure 3.**
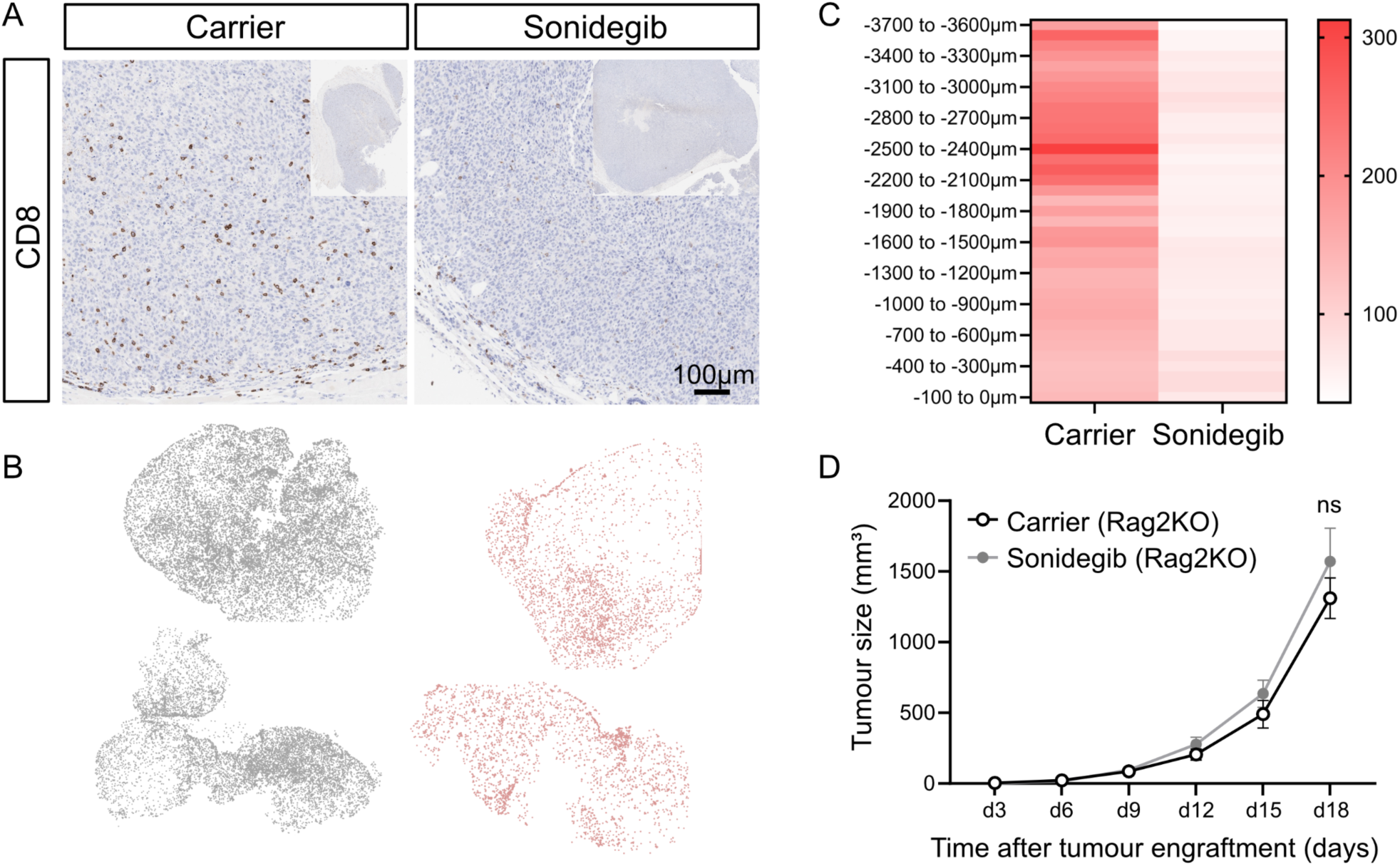
Sonidegib treatment inhibits CD8 migration into the tumour. **(A)** Representative paraffin sections from MC38 tumour-bearing mice treated with either carrier control or sonidegib and stained with an anti-mouse CD8a antibody. The whole tumour is shown in top right insert panels. **(B)** HALO analysis of CD8 cell infiltration for two representative tumours of carrier-treated (grey) and sonidegib-treated (red) mice, respectively. Each dot represents a CD8 cell. **(C)** Mean of number of CD3+CD8+ cells/ mm^2^ in 100μm-wide zones from the tumour surface (−100 to 0 μm) to the tumour centre (−3700 to −3600 μm) as quantified by HALO software (analysis workflow shown in **Suppl. Fig. 3A**). Two independent experiments, n=10 for carrier-treated and n=8 for sonidegib-treated mice. **(D)** Tumour size determined from caliper measurements. *Rag2*KO mice were subcutaneously injected with 0.5 × 10^6^ MC38 cells on d0. On d9, mice were stratified into two equal groups according to tumour size. Between d9 and d18, mice were treated daily with 20mg/kg sonidegib or carrier control by oral gavage. Two independent experiments, n=7 for carrier-treated tumours, n=7 for sonidegib-treated tumours, two-way ANOVA, mean ± SEM.

To exclude the possibility that the cytotoxic cells in tumour-bearing mice were specifically retained in secondary lymphoid tissues such as spleen and tumour-draining lymph nodes, we assessed cytotoxic cell numbers in these organs by flow cytometry. No significant change in the numbers of CD8, CD4, gammadelta T cells or NK cells was observed upon Hh inhibitor treatment in the spleen **(Suppl. Fig. 2B)**. A moderate reduction in both CD4 and CD8 T cell numbers was noted in the tumour-draining lymph node, 31% and 32%, respectively **(Suppl. Fig. 2C)**. These data suggests that the reduction of cytotoxic lymphocytes in the tumours of sonidegib-treated mice was not due to a retention of the cells in secondary lymphoid organs.

To further investigate whether the loss of tumour control **(Fig. 2D)** was due to an effect of Hh inhibitors on cytotoxic T cells or NK cells, we systemically inhibited Smo in RAG2KO mice which lack B and T lymphocytes but retain NK cells. Tumours in RAG2KO mice grow faster due to the lack of T cells, so we adopted an earlier endpoint on day 18 **(Fig. 3D)**. We observed no difference in tumour growth upon Smo inhibition indicating the exacerbated tumour growth observed in C57BL/6J mice is driven by a repressive effect of sonidegib on adaptive T lymphocytes rather than NK cells.

In summary, our data show that treating immunocompetent tumour-bearing mice with Smo inhibitors results in fewer cytotoxic T cells in the tumour with compromised ability of the cells to infiltrate deeply into the tumour mass leading to impaired tumour rejection.

### Genetic ablation of *Smo* in CD8 T cells results in reduced CD8 infiltration in the tumour and increased tumour load

To confirm that the observed lack of infiltration and tumour control in the Hh-inhibited animals was due to a direct and specific effect of the inhibitors on CD8 T cells and not due to indirect effects via other cell types or off-target effects, we generated a novel conditional knockout mouse line. In this line, *Smo* is floxed and can be inducibly deleted selectively in cytotoxic lymphocytes via an *ERT2Cre* recombinase under control of the Granzyme B promoter, circumventing any effects of *Smo* loss on T cell development. A tdTomato fluorescence reporter was included to detect Cre-mediated recombination.

MC38 tumours were subcutaneously implanted in *Smo* HET (Ctrl, *GzmB-ERT2Cre/ROSAtdTom/Smo^+/fl^*) and KO (*GzmB-ERT2Cre/ROSAtdTom/Smo^fl/fl^*) mice and excision of *Smo* in cytotoxic lymphocytes was induced by administration of Tamoxifen. Tamoxifen was given one day before tumour implantation and then again on d1, d3 and d5 post implantation **(Fig. 4A)**. *Smo* KO mice exhibited accelerated tumour growth compared to HET animals, resulting in a 2-fold increase in tumour weight at experimental endpoint **(Fig. 4B, C)**. At endpoint, the tumours in the *Smo* KO mice also harboured reduced numbers of CD8 and NK cells, albeit no statistically significant differences were observed **(Suppl. Fig. 4A)**. These experiments indicate that the effects observed in sonidegib-treated animals are reproduced by genetic ablation of *Smo* in cytotoxic lymphocytes.

**Figure 4.**
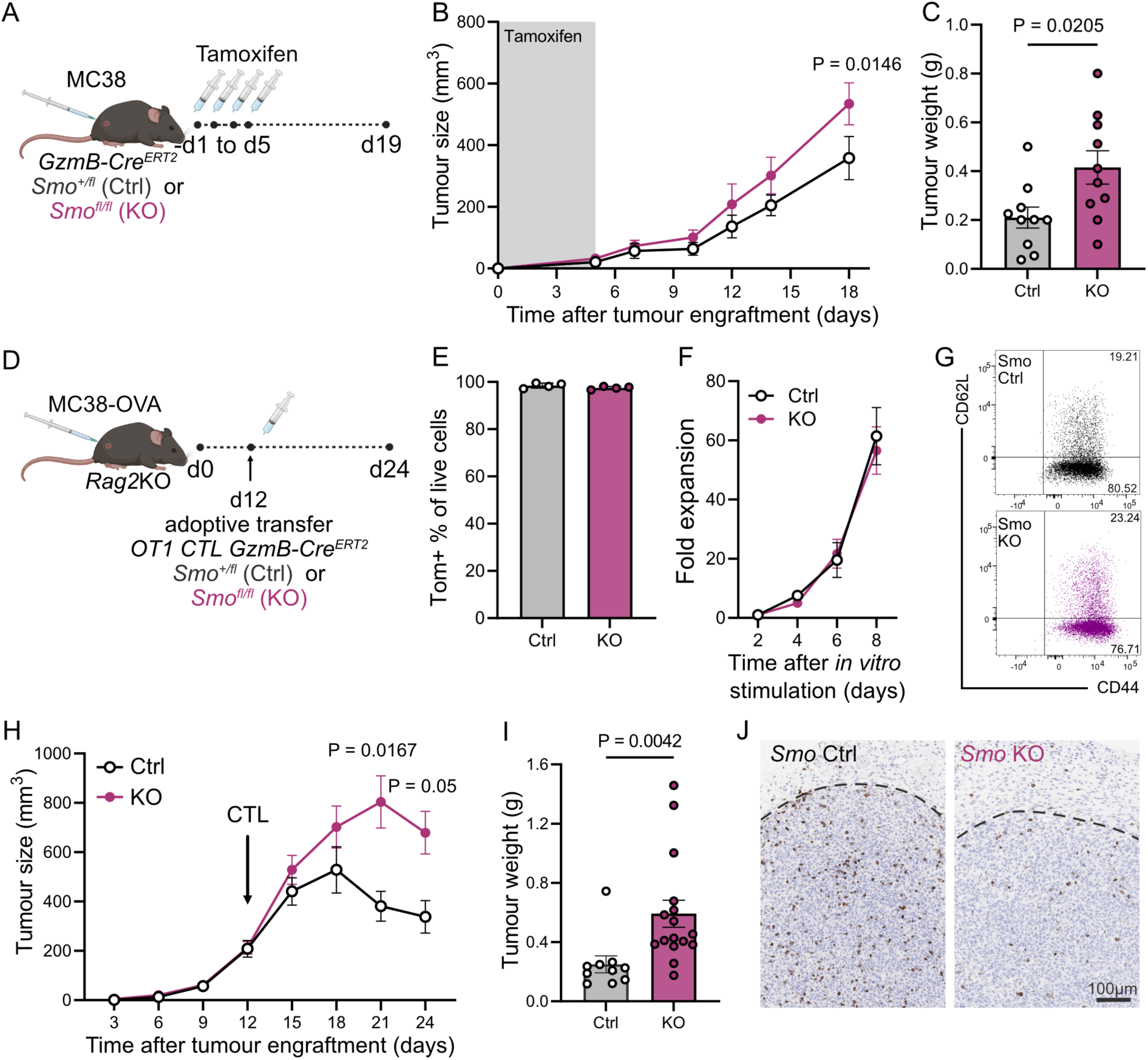
Genetic loss of *Smo* in cytotoxic T cells increases tumour growth. **(A)** Experimental Design. *GzmB-ERT2Cre/ROSAtdTom/Smo^fl/fl^*(KO) or *Smo^fl/+^* (Ctrl) animals were subcutaneously injected with 0.5 × 10^6^ MC38 cells on d0. Mice were treated with tamoxifen via intraperitoneal injections (75mg/kg) on −d1, d1, d3 and d5. **(B)** Tumour dimensions determined from caliper measurements. Four independent experiments, n=11 for *Smo^fl/+^* (Ctrl), n=10 for *Smo^fl/fl^* (KO), 2-way ANOVA, mean ± SEM. **(C)** Tumour weight at endpoint (d19). Four independent experiments, n=10 for *Smo^fl/+^* (Ctrl), n=10 for *Smo^fl/fl^* (KO), unpaired t-test, mean ± SEM. **(D)** Experimental Design. *Rag2*KO animals were subcutaneously injected with 0.5 × 10^6^ MC38-OVA cells on d0. In parallel, T cells from *GzmB-ERT2Cre/ROSAtdTom*-*OTI* mice either control or *Smo* knockout were activated with OVA peptide for 48hrs and expanded in the presence of OHT for 6-8 days. On d12, tumour-bearing mice were stratified into two equal groups according to tumour size and 1 × 10^6^ of either control (Ctrl) or Smo knockout (KO) cytotoxic T cells (CTLs) were adoptively transferred by tail vein injection. **(E)** Percentage of Tom+ cells on d7 of *ex vivo* T cell expansion, n=4 for *Smo^fl/+^* (Ctrl), n=4 for *Smo^fl/fl^* (KO), mean ± SD. **(F)** Fold expansion of CD8 cells *in vitro* following 48h stimulation with OVA peptide in the presence of OHT. Four independent experiments, n=7 for *Smo^fl/+^* (Ctrl), n=8 for *Smo^fl/fl^*(KO), mean ± SEM. **(G)** Representative flow cytometry plots of CD62L/CD44 expression on CTLs from d7 of expansion that were subsequently used for the adoptive transfers (**D, H-J** of this figure). **(H)** Tumour dimensions were determined from caliper measurements. Four independent experiments, n=10 for *Smo^fl/+^* (Ctrl), n=16 for *Smo^fl/fl^* (KO), ordinary two-way ANOVA with Geisser-Greenhouse correction and Sidak’s multiple comparison tests, mean ± SEM. **(I)** Tumour weight at endpoint (d24). Two independent experiments, n=10 for *Smo^fl/+^* (Ctrl), n=16 for *Smo^fl/fl^* (KO), unpaired Mann-Whitney test, mean ± SEM. **(J)** Representative paraffin sections from MC38-OVA tumours shown in **(H)** and stained with anti-mouse CD8α antibodies. Dotted line indicates tumour margins.

The above tumour model had the following limitations: firstly, using this schedule of Tamoxifen treatment, only an average of 40% of cytotoxic lymphocytes was excised (as determined by expression of the tdTomato fluorescent Cre reporter) **(Suppl. Fig. 4B)**; secondly, the model did not allow us to track the tumour antigen-specific T cell response; and thirdly, the GzmBER^T2^Cre also induces recombination in GzmB+ cytotoxic CD4+ T cells, gammadelta T cells and NK cells^29^. Thus, we modified our experimental design to exclusively assess the functional CD8 T cell anti-tumour response. For this purpose, we crossed our mouse line to the OTI transgenic strain to generate CD8 *Smo* KO tumour-specific CTLs *in vitro* and adoptively transferred the cells into MC38-OVA tumour-bearing, immunodeficient *Rag2KO* recipient mice. With this setup we can interrogate the role of Smo in tumour-specific T cells and without potential effects of *Smo* deletion on CTL priming, expansion, and differentiation **(Fig. 4D)**.

CD8 T cells from Ctrl and *Smo* KO mice were stimulated with OVA and treated with OH-tamoxifen *ex vivo* before being transferred into tumour-bearing mice. *In vitro*, Smo deletion was confirmed by qRT-PCR **(Suppl. Fig. 4C)** and although more than 97% of the cells were excised by d7 of culture **(Fig. 4E)**, CD8 T cell expansion or effector phenotype were not affected **(Fig. 4F, G)**. *In vivo*, Genetic ablation of *Smo* in tumour-specific CD8 T cells lead to a doubling of the tumour burden **(Fig. 4H, I)** and diminished T cell infiltration into the tumour mass **(Fig. 4J),** recapitulating the effect of sonidegib treatment.

### Genetic ablation of *Ihh* and *Gli1* in CD8 T cells does not result in reduced tumour infiltration or increased tumour load

Smo is the main signal transducer of the Hh pathway and the molecular target of almost all clinically used inhibitors, including vismodegib and sonidegib. To investigate mechanistically whether the migration phenotype we observed was perhaps regulated by Hh ligands upstream or the Gli transcription factors downstream of Smo **(Fig. 2A)**, we assessed CD8 T cells deficient in either *Indian Hh* (*Ihh*) or *Gli1*. Previous work from our laboratory has shown CD8 CTLs only express Ihh as a Hh ligand (and not Dhh or Shh)^21^ and T cells are unable to respond to exogenous Hh ligands^21, 30^. Furthermore, Gli1 is the main transcription factor expressed in CD8 CTLs^21^.

OTI-CTLs deficient in *Ihh* or *Gli1* were generated *ex vivo* from our respective mouse models and adoptively transferred into MC38-OVA tumour-bearing *Rag2KO* recipient mice. Loss of *Ihh* or *Gli1* in CD8 T cells did not alter proliferation or effector phenotype *in vitro* **(Suppl. Fig. 4F, G, L, M)**.

*In vivo*, OTI-CTLs deficient in *Ihh* or *Gli1 were* equally effective as their wildtype counterparts at controlling tumour growth **(Suppl. Fig. 4H, N)**. While tumour weights were identical in the mice treated with *Ihh* Ctrl versus KO OTI-CTLs, we observed a reduced, albeit non-significant, ability of *Gli1*KO OTI-CTLs to control MC38-OVA tumours *in vivo* **(Suppl. Fig. 4N, O)**. Furthermore, the levels of tumour infiltration by CD8 cells were unaffected by the loss of either *Ihh* or *Gli1* in OTI-CTLs **(Suppl. Fig. 4J, P)**.

These data show that, unlike genetic ablation of *Smo*, loss of *Ihh* or *Gli1* in cytotoxic CD8 T cells, does not affect tumour growth or restrict CD8 migration into the tumour microenvironment in the models tested.

### Pharmacological inhibition or genetic ablation of *Smo*, but not *Ihh* or *Gli1*, inhibits CTL migration towards tumour cells

To investigate whether the diminished MC38 tumour infiltration by CD8 T cells seen upon sonidegib treatment and genetic ablation of *Smo* in the T cells was indeed caused by reduced migratory capacity, we utilised a transwell migration assay. Here, MC38-OVA cells were plated in the bottom well and OTI-CTLs were added to the insert **(Fig. 5A)**.

**Figure 5:**
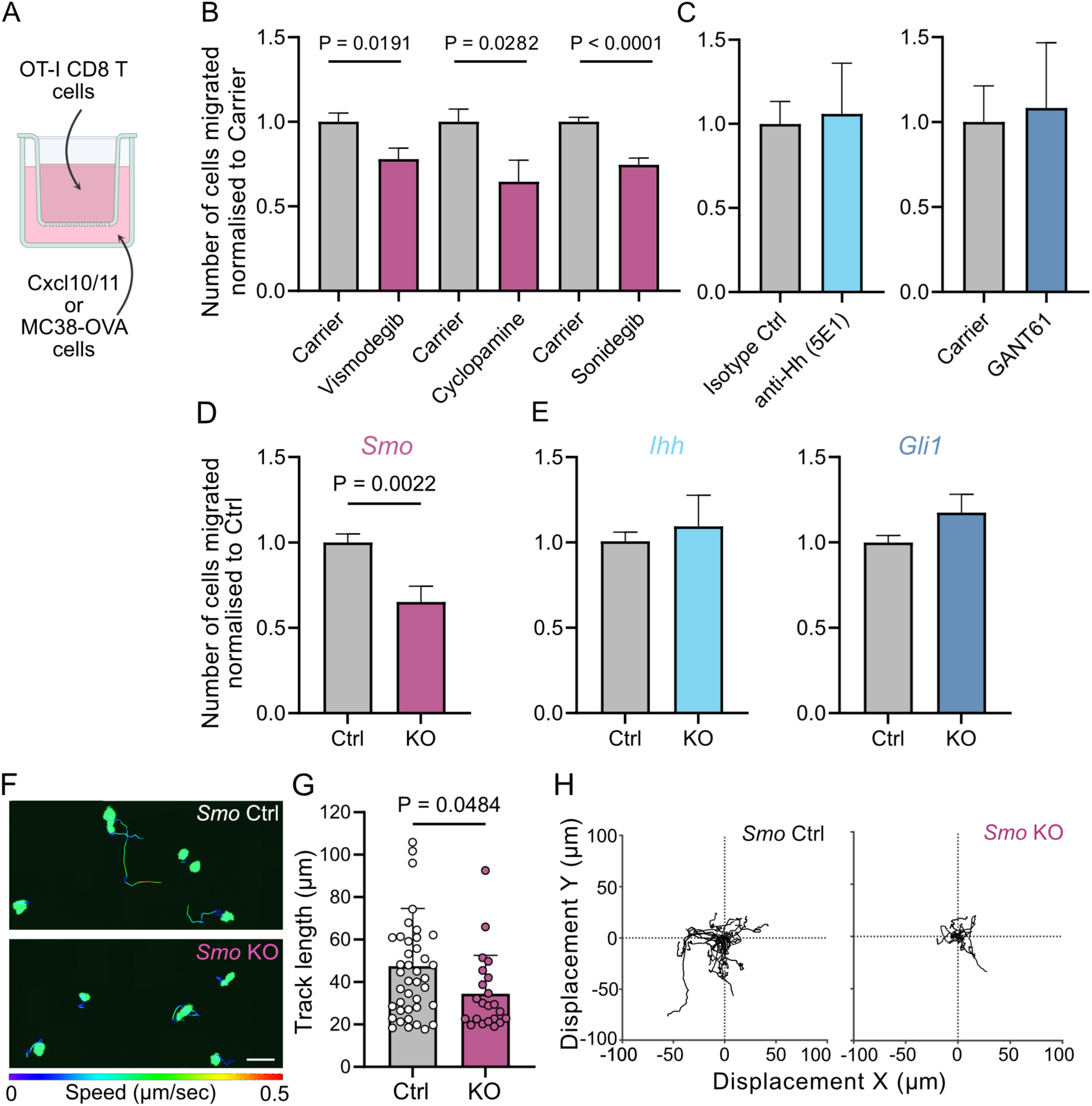
Loss of *Smo*, but not *Ihh* or *Gli1*, inhibits CD8 migration *in vitro*. **(A)** Schematic of transwell experiment to assess directed migration. Cxcl10 and Cxcl11-supplemented T cell media **(B, C)** or MC38-OVA cells **(D, E)** were supplied in the bottom well. A 5μm transwell insert was placed on top and CTLs on d6-8 of culture were dispensed in the insert. After 6hrs, cells from the bottom well were collected for flowcytometric analysis. **(B)** T cell migration in the presence of Smo inhibitors. Vismodegib, n=6, two independent experiments; Cyclopamine, n=6, two independent experiments, and Sonidegib, n=8, two independent experiments. For each drug, paired t-tests were performed, and mean ± SEM is shown. **(C)** T cell migration in the presence of Hh ligand blocking antibody 5E1 and Gli inhibitor GANT61. 5E1: n=4, two independent experiments, GANT61: n=3, two independent experiments. For each drug, paired t-tests were performed, and mean ± SEM is shown. **(D)** CTLs were generated from conditional *Smo^fl/+^or Smo^+/+^* (Ctrl) as well as *Smo^fl/fl^* (KO) mice and migratory capacity was assessed. Migration is normalised to Ctrl. n=4 for Ctrl, n=4 for KO. All samples were run in technical triplicates across three independent experiments. Unpaired t-test, mean ± SEM. **(E)** CTLs were generated from conditional *Ihh^fl/+^* (Ctrl) or *Ihh^fl/fl^* (KO) mice as well as *Gli1^+/+^* (Ctrl) or *Gli1^eGFP/eGFP^*(KO) mice and migratory capacity was assessed. Migration is normalised to Ctrl. n=4 for *Ihh* Ctrl, n=5 for *Ihh* KO, n=5 for *Gli1* Ctrl and n=5 for *Gli1* KO. All samples were run in technical triplicates across three to four independent experiments. Unpaired t-test, mean ± SEM. **(F-H)** CTLs were generated from conditional *Smo^fl/+^or Smo^+/+^* (Ctrl) as well as *Smo^fl/fl^* (KO) mice and migratory capacity of cells was assessed by live imaging on Icam-coated glass slides for 20min. **(F)** Representative snapshots of live imaging with individual cell tracks shown. Track gradient denotes speed (blue to red, 0 to 0.5 μm/sec) and scale bar equals 20 microns. **(G)** Mean track length of CFSE-labelled *Smo* Ctrl and KO CTLs is shown. Each dot represents an individual cell. n=3 *Smo* Ctrl and n=3 *Smo* KO, one independent experiment. Unpaired t-test, mean ± SD. **(H)** XY displacements graphs for representative samples shown in **(F)**.

Addition of Smo inhibitors, including the clinically approved vismodegib and sonidegib as well as the *bona fide* Smo inhibitor cyclopamine reduced CTL migration by 20-40% **(Fig. 5B)**. By contrast, treating the CTLs with a 5E1 antibody that blocks Hh ligands (Ihh, Shh, Dhh) from binding to Ptch1, or the Gli1 antagonist GANT61, had no effect on CTL migration **(Fig. 5C)**. All inhibitors tested did not affect T cell viability **(Suppl. Fig. 5K)**.

Taken together the data is consistent with our *in vivo* findings using *Ihh* and *Gli1* KO T cells **(Suppl. Fig. 4E-P)**, suggesting the migration effect is mediated by Smo.

To confirm that the reduced migration observed in the presence of Smo inhibitors was indeed Smo-specific and not caused by toxicities, we performed transwell assays using *Smo* KO OTI-CTLs. We observed a 35% reduction of the ability of *Smo* KO CTLs to reach the tumour cells compared to *Smo* Ctrl CTLs **(Fig. 5D)**. Genetic deletion of either *Ihh* or *Gli1* again did not affect the CTL migration in the *in vitro* system **(Fig. 5E)**.

To further investigate Smo-dependent CTL migratory capacity in real-time, we developed a live cell imaging system. Here, we use a spinning disk confocal microscope to track CTL migration on ICAM-coated glass over 20 minutes. *Smo* KO CTLs displayed reduced speed **(Fig. 5F)** with overall 27% reduced track length **(Fig. 5G)** resulting in lower displacement from their starting point compared to *Smo* WT cells **(Fig. 5H)**.

Our *in vitro* migration assays reveal that Smo, but not canonical Hh signaling through Ihh or Gli1, is critical for the migration of CTLs towards tumour cells.

### Smo regulates CTL migration via its GPCR function

We have shown that Smo is critical for CTL migration *in vitro* and *in vivo* and acts independently of both the Hh ligand Ihh and the transcription factor Gli1. T cell migration into the tumour microenvironment is guided by chemokine gradients. A small set of chemokines are secreted by tumour cells and are recognised by chemokine receptors expressed on T cells^31^.

We were interested to investigate whether Smo might broadly regulate the expression of chemoattractants and receptors in T cells. We first profiled cytokine and chemokine expression in *Smo* WT and KO CTLs using a XL-cytokine and chemokine array **(Suppl. Fig. 5A, B)**. Out of 111 targets probed, only few were expressed in CTLs **(Suppl. Fig. 5A)**, and quantification of the cytokines and chemokines expressed did not show significant differences between *Smo* wildtype and *Smo* KO cells **(Suppl. Fig. 5B)**. In addition, mRNA expression of the top hits from the array was also not significantly altered upon loss of *Smo* **(Suppl. Fig. 5C)**.

We next determined the main chemokines expressed by the tumour cells used in our *in vitro* migration assays and the tumour-bearing mice models. MC38-OVA cells express high levels of Cxcl10 and compared to Cxcl10, 87-fold lower levels of Cxcl11 and 3732-fold lower Cxcl9. Ccr1, 3, 5, Cxcl12 and Cxcr4 were barely detected **(Suppl. Fig. 5D)**. All three expressed cytokines, Cxcl9, 10, and 11 bind to the Cxcr3 receptor. Thus, we profiled Cxcr3 expression in Ctrl or *Smo* KO CTLs by qRT-PCR **(Suppl. Fig. 5E),** flow cytometry and immunoblotting **(Suppl. Fig. 5F, G)**. The loss of *Smo* did not lead to any significant alterations in Cxcr3 expression. To investigate whether the Cxcr3-Cxcl9/10/11 signaling axis is active in the transwell migration assay, we treated CTLs with either the Cxcr3 antagonist SCH546738 or a blocking antibody against Cxcr3 (α-Cxcr3). Both treatments resulted in a reduction of CTL migration by 31% and 20%, respectively **(Suppl. Fig. 5H)**. However, no additive effect on migration was observed when *Smo* Ctrl and KO cells were treated with SCH546738 or α-Cxcr3, even when increasing concentrations of the inhibitor were used **(Suppl. Fig. 5I, J)**. Taken together, these data suggest that genetic ablation of *Smo* in CTLs has no effect on the CTL chemokine profile and although CTLs still utilise the Cxcr3-Cxcl9/10/11 axis for migration, this dependency is not differentially regulated in *Smo* KO CTLs.

Apart from being the key signal transducer for canonical Hh signaling, Smo has been shown to function as a class F G-protein coupled receptor (GPCR) (reviewed in^32^) **(Fig. 6A)**. Importantly, previous work in fibroblasts has shown that Smo can regulate migration via its GPCR function^33^. To determine whether a similar effect may be at play in CTLs, we blocked coupling of Smo to G_i_ subunits by treating the cells with *Pertussis Toxin* (PTX). PTX blocks G protein-GPCR interactions by catalysing the ADP-ribosylation of α_i_ subunits^34^. Notably, PTX led to a 74% reduction in CTL migratory capacity **(Fig. 6B, Suppl. Fig. 6A)**.

**Figure 6:**
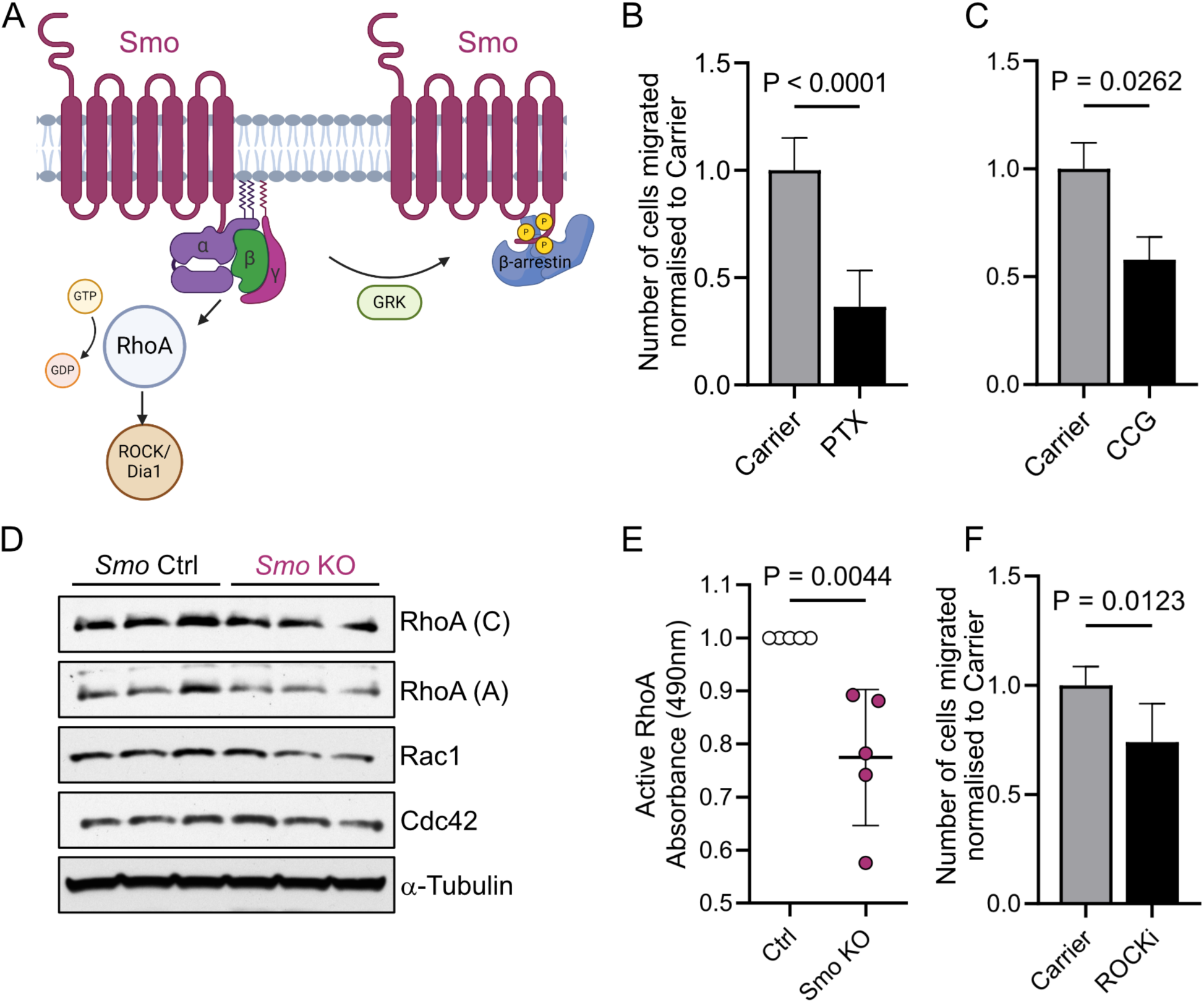
Genetic loss of *Smo* affects cytotoxic CD8 migration via GPCR coupling. **(A)** Smoothened (Smo) is a class F 7-TM G-protein coupled receptor. Smo has been shown to utilise Gαi heterotrimeric proteins and regulate RhoA and Rac1 activation. After GPCR activation, GRKs regulate the phosphorylation of the Smo C-terminus and recruitment of arrestins^32^. **(B, C)** Transwell migration assays with Cxcl10 and Cxcl11-supplemented T cell media at the bottom well. CTLs on d6-8 of culture in drug-containing media were dispensed in the insert. After 6hrs, cells from the bottom well were collected for flow cytometric analysis. *Pertussis* toxin (PTX) 100ng/ml, five independent experiments, n=13. CCG215022 0.32mM, two independent experiments, n=6. All samples were run in technical quadruplicates. For each drug paired t-tests were performed and mean ± SD is shown. **(D, E)** CTLs were generated from conditional *Smo^fl/+^or Smo^+/+^*(Ctrl) as well as *Smo^fl/fl^* (KO) mice. **(D)** Western blot analysis of *Smo^fl/+^* (Ctrl) and *Smo^fl/fl^* (KO) CTL lysates. Two separate antibodies were tested for RhoA, one form Cytoskeleton (C) and one from Abcam (A). One independent experiment, n=3 for *Smo* Ctrl and n=3 for *Smo* KO, quantification can be found in **Suppl. Fig. 6B**. **(E)** Active RhoA G-lisa. Three independent experiments, n=5 for Smo Ctrl and n=5 for Smo KO, unpaired t-test, mean ± SD. **(F)** Transwell migration assays as in **(B, C)**. ROCK inhibitor 1 μM, two independent experiments, n=5, paired t-test mean ± SD.

Other studies have suggested that after Smo activation, the Smo c-tail can become phosphorylated, leading to an altered conformation permissive to β-arrestin recruitment. Recruitment of β-arrestin to the active conformation can then drive high-level GPCR signal propagation and eventual Smo desensitization^32^. To determine if c-tail phosphorylation and arrestin recruitment could play a role in CTL migration, we used a G protein-coupled receptor kinase (GRK) inhibitor, CCG215022, to block kinase activity, inhibiting GPCR phosphorylation. Treating CTLs with the GRK inhibitor resulted in a 43% decrease in CTL migratory capacity **(Fig. 6C, Suppl. Fig. 6A)**.

To understand the mediators of T cell migration downstream of Smo GPCR activity, we turned to RhoA, a small GTPase which has been shown to regulate fibroblast migration^33^. First, we confirmed expression of RhoA and other small GTPases including Rac1 and Cdc42 at steady state in CTLs by bulk RNAseq **(Suppl. Fig. 6B)**. We then compared protein levels of total RhoA, Cdc42, and Rac1 in *Smo* Ctrl and KO CTLs which were not significantly altered **(Fig. 6D, Suppl. Fig. 6C)**. However, expression of RhoA and Rac1 were marginally reduced in the KO.

Next, we assessed the effect of *Smo* loss on active RhoA. Intriguingly, levels of GTP-bound, active RhoA in *Smo* KO CTLs were reduced by 22% **(Fig. 6E)**. In addition, CTLs treated with a Rho kinase inhibitor, displayed 26% reduced migration compared to vehicle-treated controls **(Fig. 6F, Suppl. Fig. 6A)**.

This suggests that CTLs rely on GPCR coupling, RhoA and c-tail phosphorylation to migrate. CTLs with genetic ablation of *Smo* have lower levels of active RhoA, resulting in diminished migratory capacity.

### Vismodegib treatment reduces T cell infiltration in basal cell carcinoma

To determine whether human CTLs also rely on SMO to migrate, we performed transwell assays with SMO inhibitors vismodegib and sonidegib. The migratory capacity of human CD8 T cells isolated from healthy donors was reduced in the presence of vismodegib by 14% and sonidegib by 21% **(Fig. 7A).** Both inhibitors did not affect T cell viability at the concentrations used **(Suppl. Fig. 7A)**.

**Figure 7:**
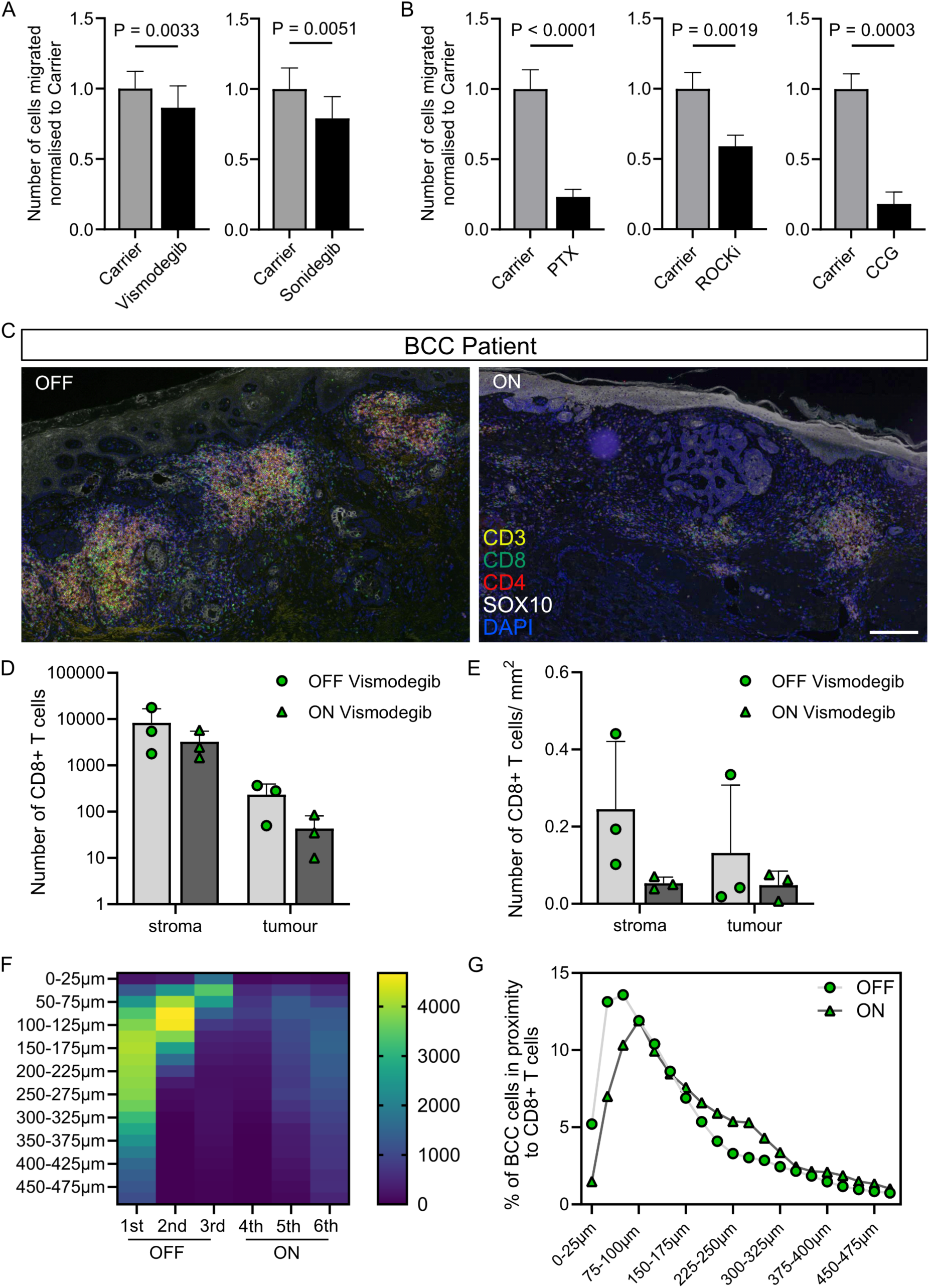
Inhibition of SMO diminishes human CD8 T cell migration *in vitro* and in the clinic. **(A)** Transwell assays with CXCL10 and CXCL11-supplemented T cell media at the bottom well. Inserts coated with matrigel were placed on top of the well and d10-12 CTLs were dispensed in the insert in drug-containing media. Vismodegib 5 μM, one independent experiment, n=5 healthy donors. Sonidegib 10 μM, one independent experiment, n=5 healthy donors. For each drug paired t-tests were performed and mean ± SD is shown. **(B)** Transwell assays as in **(A)**. *Bortedella pertussis* toxin (PTX) 200ng/ml, two independent experiments, n=9 healthy donors. ROCK inhibitor 10 μM, two independent experiments, n=6 healthy donors, GRK inhibitor CCG215022 0.032-0.32mM, three independent experiments, n=11 healthy donors. For each drug paired t-tests were performed and mean ± SEM is shown. **(C)** Representative sections of separate BCCs resected from the same cancer patient while *“on”* or *“off”* vismodegib treatment and stained with antibodies against CD3, CD8, CD4, SOX10. Nuclei were stained with DAPI. Vismodegib treatment diminished the number of T cells infiltrating the tumour. Scale bar equals 100 μm. **(D)** Numbers of CD8 T cells found in the stroma (defined as within 500 μm outwards of the tumour margin) or the tumour (defined as within the tumour margins). Separate resected tumour specimens of the same patient (ALF006) spanning 11 years while the patient was *“on”* or *“off”* vismodegib treatment were analysed. n=3 *“on”* and n=3 *“off”* treatment, mean ± SD is shown. **(E)** Numbers of CD8 T cells found in the stroma (defined as within 500 μm outwards of the tumour margin) or the tumour (defined as within the tumour margins) per mm^2^ of stroma and tumour areas, respectively. n=3 *“on”* and n=3 *“off”* treatment, mean ± SD is shown. **(F)** Proximity analysis of each BCC cell to its closest neighbouring CD8 T cell (analysis workflow shown in **Supp. Fig. 7C-E**). Heatmap indicates the number of BCC tumour cells within the indicated distance brackets. n=3 BCCs *“on”* and n=3 BCCs *“off”* treatment. **(G)** Percentage of total BCC cells shown with regards to their closest neighbouring CD8 T cell (analysis workflow shown in **Supp. Fig. 7C-E**) adjusted for tumour load. Means of n=3 BCCs *“on”* and n=3 BCCs *“off”* treatment are shown.

We then treated human CD8 T cells with PTX, Rho kinase and GRK inhibitors, to establish whether, like in murine CD8 T cells, the GPCR machinery was driving human CTL migration. Human T cells were even more sensitive to all three drugs compared to murine T cells, leading to a reduction in migratory capacity by 77% (PTX), 41% (ROCKi) and 82% (GRKi), respectively **(Fig. 7B)**. The viability of primary human T cells was unaffected by inhibitor treatment **(Suppl. Fig. 7B)**.

In order to assess if our findings apply to patients treated with SMO inhibitors in the clinic, we studied cancer patients with multiple Basal Cell Carcinomas (BCCs). A small subset of patients with locally advanced or metastatic BCCs who cannot be treated with surgery or radiation are systemically treated with the SMO inhibitor vismodegib in the UK. We initiated a clinical research study at Addenbrooke’s Hospital (<u>A</u>ltered <u>L</u>ymphocyte <u>F</u>unction in health and disease, ALF study) and recruited 11 BCC patients treated with vismodegib in an *“on”*/*“off”* treatment cycle pattern. We sought to determine whether the number of CD8 T cells in BCCs was affected by treatment with vismodegib. For 5 of the BCC patients, we were able to collect rare tumour biopsies when patients are either *“on”* or *“off”* treatment with vismodegib **(Suppl. Fig. 7F)**.

Whilst the sample pool is limited, for one of our patients we obtained six separate tumour resections, either excised when the patient was *“on”* treatment with vismodegib (ON) or during treatment break (OFF) over a period of 11 years **(Fig. 7C-G, Suppl. Fig. 7G-I)**. As observed in our murine cancer models, infiltration of CD8 T cells into the TME was reduced upon treatment with SMO inhibitor shown by immunofluorescence **(Fig. 7C)**. This was true for total numbers of CD8 T cells in the tumour and stroma **(Fig. 7D)** and for CD8 T cell counts adjusted to surface area **(Fig. 7E)**. Vismodegib treatment did not lead to a reduction of CD4 T cells in the TME **(Suppl. Fig. 7G)**.

CD8 T cells need to be in close/direct contact to tumour target cells to efficiently carry out their function^35^. Thus, we next calculated the distance between each BCC cell and its nearest CD8 T cell (workflow shown in **Suppl. Fig. 7C-E**). We observed that more BCC tumour cells were near CD8 T cells when patients were *“off”* Hh inhibitor treatment **(Fig. 7F, Suppl. Fig. 7F)**. Importantly, when assessing adjacent cells (intercell distance less than 50 μm), 18% of BCC cells were neighbouring CD8 T cells when patients were *“off”*, compared to 8% when *“on”* vismodegib treatment **(Fig. 7G)**. The number of BCC cells near CD4 T cells was also slightly enriched when the patient was “*off*” treatment **(Suppl. Fig. 7H, I).**

Taken together, our data indicate that vismodegib treatment has a previously unappreciated effect on the CD8 T cell response, inhibiting T cell migration. This effect is distinct from the direct anti-tumour effect of vismodegib on the cancer cells themselves. Together, our work might explain why vismodegib did not have the desired efficacy across cancer types in the clinic.

## DISCUSSION

Potent inhibitors of the Hh pathway, mainly targeting the key signal transducer SMO, have been developed and shown efficacy in BCC^36^ and SHH medulloblastoma^37^ which harbour Hh driver mutations. However, Hedgehog inhibitors have repeatedly failed to meet primary endpoints in clinical trials for cancers where Hh signaling is aberrantly amplified **(Fig. 1, Suppl. Fig. 1)**. Especially poor responses have been observed in the clinic when Hh inhibitors have been used for the treatment of gastrointestinal cancers.

Guided by the clinical data, we chose a murine model of colorectal cancer, where Hh inhibitors performed poorly. Strikingly, we observed that Hh inhibition with sonidegib (LDE225) treatment led to a doubling of the tumour burden in the animals. Of note, we have shown that sonidegib treatment also did not reduce the tumour burden in a subcutaneous KPC-derived pancreatic cancer model (2838c3) or a subcutaneous melanoma model (Yummer 1.7) **(Suppl. Fig. 8)**. This is in contrast to published work in a murine model of PDAC using the same inhibitor which resulted in a reduction of tumour burden^38^. The discrepancy in tumour response is likely due to the different model system used with regards to the tumour cell line and site of implantation.

Unlike previously published work, Hh inhibition in our model had no effect on the TME overall where tumour stroma, blood vessels and main immune subsets including macrophages and B cells remained unchanged. Previous work in murine models of breast cancer had observed a shift in macrophage polarisation from M1 to M2 upon vismodegib treatment with no effect on tumour growth reported^39^. We did not observe a change in overall macrophage abundance by F4/80 staining or M1/M2 polarisation using ConsensusTME.

Instead of broad changes in the TME, we find that Hh inhibition specifically abolishes cytotoxic lymphocytes infiltration into the tumour - especially the infiltration of cytotoxic CD8 T cells that are essential for tumour control. We further characterised the migration defect using bespoke *Smo* knockout models and adoptive transfer experiments and confirmed that the migration defect is T cell intrinsic. Migration assays and live imaging approaches revealed that CTLs lacking *Smo,* or ones treated with range of clinical and non-clinical Smo inhibitors have profoundly diminished migratory capacity.

Additional work using knockout models of *Ihh* and *Gli1* as well as inhibitors and agonists of the pathway, demonstrated that CTL migration relies on non-canonical Hh signaling via Smo, independently of exogenous and endogenous Hedgehog ligands or a Gli-mediated transcriptional programme. Furthermore, the reduced migratory capacity could not be attributed to altered expression of chemokine receptors on the cell surface of CTLs. Instead, we find that it is the GPCR function of Smo that regulates migration and that Smo knockout CTLs have reduced levels of active, GTP-bound RhoA.

A role for the GPCR function of Smo via RhoA has been previously described in fibroblast migration^33^. In that paper, migration is linked to both G_i_ coupling and PI3K activity downstream of Smo, but independent of the Smo c-tail^33^. The c-tail of Smo is necessary for Gli activation and β-arrestin recruitment but dispensable for Gi coupling^40^. In our work, we have established a role for G_i_ coupling (PTX) and RhoA (ROCKi) in CTL migration similarly to what has been described for fibroblasts, but we have also established a role for β-arrestin recruitment (CCG). In the future it would be interesting to determine the contribution levels of Smo versus other GPCRs towards G_i_ coupling, RhoA activation, and β-arrestin recruitment during CTL migration.

Finally, we asked whether human CD8 T cells would also rely on the GPCR function of SMO to migrate. For this, we treated CD8 T cells isolated from healthy donors with SMO inhibitors vismodegib and sonidegib as well as GPCR inhibitors to show that both SMO and the GPCR machinery are critical for the migration of human T cells.

To investigate whether Hh inhibitor treatment in cancer patients would also affect T cell migration towards the tumour mass, we recruited BCC patients and obtained resected tumour specimens. We collected 6 matched samples (3 *“on”* and 3 *“off”* vismodegib) from the same patient obviating a confounding effect of different immune systems from different patients. CD8 T cells in the tumour stroma and within the tumour cell nests were reduced upon vismodegib treatment as was the percentage of tumour cells near CD8 T cells.

Here, we provide the first comprehensive functional assessment of the anti-tumour immune response upon Hh inhibition *in vivo* and reveal a profound and selective effect on cytotoxic T cells infiltration while other immune subsets are less affected.

Our work has substantial implications for the clinic and potentially explains why trials of Hedgehog inhibitors in solid tumours have yielded mixed results. Furthermore, it might provide a roadmap for an improved Hh inhibitor trial design which would selectively target the tumour but spare the immune compartment.

One approach would be to ensure sufficient treatment breaks when patients are on SMO inhibitors to allow T cell infiltration into the tumour microenvironment. This could be extremely important when Hh inhibition is combined with immune checkpoint blockade such as anti-PD1 (Pembrolizumab). Another approach would be to selectively target Hh ligands and their processing, e.g. through Hedgehog acetyltransferase inhibitors^41, 42^, and thus avoiding inhibitory effects on CD8 T cell migration.

Most importantly, the work uncovers Hh as a novel regulator of cytotoxic T cell migration into the tumour microenvironment which might inform new strategies to increase migration in T cell therapeutics.

## ONLINE METHODS

### Mice

*RAG2*KO were a generous gift from Suzanne Turner (University of Cambridge) and OTI mice were purchased from the Jackson Laboratory (*C57BL/6-Tg (TcraTcrb*)1100Mjb/j, Stock no. 003831). OTI *RAG2*KO mice were generated from these. *Smo^f/f^* (Stock no. *Smo^tm2Amc^/J*, 004526) and *Ihh^f/+^* (*Ihh^tm1Blan^*/J, Stock no. 024327) were purchased from The Jackson Laboratory. *GzmB^ERT2Cre^/ROSA26EYFP* mice^29^ were a generous gift from D. T. Fearon (Cold Spring Harbor Laboratory). *dLckCre* and *ROSA26tdTom* mice were a generous gift from Randall Johnson and Douglas Winton (University of Cambridge), respectively. Smo^fl/fl^ mice were crossed to *GzmB^ERT2Cre^/ROSA26tdTom* mice to generate *GzmB^ERT2Cre^/ROSAtdTom/Smo^fl/fl^*mice. *Ihh^fl/fl^* mice were crossed onto *dLckCre/ ROSA26tdTom* mice. *Gli1-eGFP* mice^43^ were a generous gift from Alexandra Joyner (Sloan Kettering Institute). All strains were backcrossed onto the C57BL/6J background (Charles River, UK) for more than 10 generations.

Mice were genotyped using Transnetyx, maintained at the CRUK Cambridge Institute-University of Cambridge and used after 6 weeks of age. All housing and procedures were performed in strict accordance with the United Kingdom Home Office Regulations.

### Tumour models

C57BL/6J or *Rag2*KO or *GzmB-Cre^ERT2^Smo^+/fl, fl/fl^* mice were injected subcutaneously with tumour cells on d0. MC38 (Kerafast #ENH204-FP) or MC38-OVA cells were injected at 0.5 × 10^6^ in PBS. MC38-OVA were generated by retroviral transduction of MC38 cells with HEK293T supernatant transfected with a pMig-cytoOVA-IRES-tdTomato plasmid. For *GzmB-Cre^ERT2^Smo^+/fl, fl/fl^* adoptive transfer, mice were treated with 75mg/kg tamoxifen (Merck #T5648) diluted in corn oil (Merck #C8267) one day before tumour implantation (-d1) and then again on d1, d3 and d5. For KPCY line 2838c3^44^ (Kerafast #EUP013-FP) cells were co-injected 1:1 with Matrigel (Cultrex, Bio-techne #3433-010-R1). YUMMER 1.7 melanoma cells were a generous gift from Marcus Bosenberg^45^ and were injected at 0.5 × 10^6^ in PBS. All tumour lines tested negative for mouse pathogens (M1 PCR panel, Surrey Diagnostics). Tumours were grown for a total of 16-24 days, as indicated in respective experimental designs. Mice were treated with sonidegib (sonidegib diphosphate salt (LC labs #S-4699) dissolved in polyethylene glycol 400 (Sigma #91893) and 5% dextrose (Sigma #G8270) in water, 75:25 v/v) or carrier control and administered by oral gavage at 20mg/kg daily for up to 12 days. For adoptive transfer experiments of cytotoxic CD8 T cells, mice were first stratified into two equal groups according to tumour size, and then 1 × 10^6^ cells were injected intravenously on d12. Tumour size was assessed by caliper measurements and tumour volume was calculated using the formula:

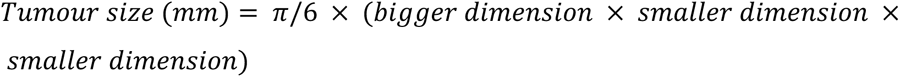

### Tissue culture and cell lines

Murine CD8 T cells were cultured at 1×10^6^ cells/ml in complete RPMI media: RPMI 1640 Medium (Thermo #21875034) supplemented with 10% heat-inactivated, batch-tested FCS (Biosera #1001), 50µM β-Mercaptoethanol (50mM, Thermo #31350010), 100U/ml Penicillin/Streptomycin (10,000 U/ml, Thermo #15140122), 1µM Sodium Pyruvate (Gibco, #11360039), 10µM HEPES (Sigma #H0887) and 10ng/ml murine IL-2 (Peprotech #212-12).

HEK293T, MC38, MC38-tdTom, MC38-Cyto-OVA-tdTom and KPCY (clone 2838c3) cells were cultured in 10% heat-inactivated FCS DMEM (Thermo #41966029). YUMMER 1.7-tdTom were cultured in 10% FCS Ham’s F-12 Nutrient Mix (Thermo #21765029) supplemented with NEAA (Thermo #11140035).

All cells were grown in a humidified incubator at 37°C and 5% CO2. Cell lines were STR profiled and all tested mycoplasma negative (PhoenixDx mycoplasma qPCR kit, Procomcure #PCCSKU15209).

### Murine CD8 cell isolation and expansion and *in vitro* treatment

Total murine CD8 T cells were isolated using MACS (negative selection, Miltenyi Biotec #130-104-075) according to the manufacturer’s instructions. The purity of the sorted populations was above 95%. CD8 T cells were stimulated with platebound anti-CD3ε (1µg/ml, eBioscience #16-0033-86) and anti-CD28 (2µg/ml, eBioscience #16-0281-86) antibody. Alternatively, cell suspensions from spleen and lymph node from OTI mice were stimulated with 10nM Ova257-264 peptide (SIINFEKL, Cambridge Bioscience #60193-5-ANA). T cells were stimulated for 48hrs and grown for up to 11 days. Cells were considered cytotoxic (CTL) between d6-d8 and used at that time for adoptive transfers, killing assays and migration assays. Cells were restimulated on d10 with 1μg/ml plate-bound anti-CD3e (eBio500A2 (500A2), functional grade, eBioscience #16-0033-86) for creation of cell pellets for RNA or protein extraction.

For CTL generation from *GzmBCre^ERT2^/ROSAtdTom/Smo^+/fl,fl/fl^* mice *in vitro*, cells were incubated with 4-hydroxytamoxifen (4-OHT, Bio-techne #341210) diluted in DMSO (Merck #D12345) in order to activate the CreERT2 recombinase. Cells were treated at a concentration of 300nM and fresh 4-OHT was added daily for the first five days of culture. Control-treated cells received DMSO in complete RPMI.

### Human CD8 cell isolation and expansion and *in vitro* treatment

PBMCs were obtained from peripheral blood samples. Samples were taken from sporadic BCC patients enrolled in the ALF research study (Research into <u>A</u>ltered <u>L</u>ymphocyte <u>F</u>unction in Health and Disease, IRAS 220302, 17/YH/0304). Buffy coats were obtained from NHS Blood and Transplant under the ALF ethics.

To isolate PBMCs, total blood products were diluted in 2% FCS PBS and laid over Ficoll density gradient medium (Merck #17-1440-02) in SepMate 50ml tubes (Stemcell Technologies, #85460). Subsequently, CD8 cells were isolated using the MojoSort™ Human CD8 T Cell Isolation Kit (Biolegend # 480129) according to the manufacturer’s instructions. Cells were stimulated for 48h with ImmunoCult™ Human CD3/CD28/CD2 T Cell Activator (25 µL/mL/1×10^6^ CD8 cells; Stemcell Technologies, #10970). CD8 T cells were cultured in Immunocult (Stemcell Technologies #10981) supplemented with 100 U/ml human IL-2 (Miltenyi Biotec #130-097-746) and 100 U/ml Penicillin/Streptomycin (Gibco).

### CD8 T cell migration assay

Incucyte chemotaxis and reservoir plates were used (Clear View 96-well Chemotaxis plate for cell migration and reservoirs, Sartorius #4582 and #4600). Insert wells were coated with 0.5% Matrigel (Cultrex, Bio-techne #3433-010-R1) in PBS.

CTLs were pre-treated with small molecules or carrier controls at the indicated concentrations (Table 1) for 1hr in T cell media before the assay. 60μl of cell suspension containing small molecules were plated in each well of the insert plate containing 7000 CTLs/well. The reservoir plate was filled with pre-warmed media containing 200nM Cxcl10 and 125nM Cxcl11 or seeded tumour cells. The insert plate was placed on top of the reservoir plate and left for 6-8hrs at 37°C and 5% CO_2_. Afterwards, the insert plate was discarded and cells from the reservoir plate were aspirated, stained, and enumerated by flow cytometry.

**Table 1:** Small molecules used for migration assays.

| Smal molecule | Concentration | Carrier | Cat no. | Supplier |
| --- | --- | --- | --- | --- |
| Vismodegib | 5µM | DMSO | V-4050 | LC Labs |
| Cyclopamine | 5µM | ETOH | J61528 | Thermo |
| Sonidegib | 10µM | PEG/Dextrose/Water | S-4699 | LC Labs |
| Anti-Hh ligand<br>5E1 | 10µg/ml | PBS | Ab01175-23-0 | 2Bscientific |
| GANT61 | 5µM | ETOH | 1892-5 | Cambridge Bioscience |
| SCH546738 | 100nM-1µM | DMSO | HY-10017 | Cambridge Bioscience |
| Anti-Cxcr3 | 200 ng/ml | 0.09% NaN<sub>3</sub> in PBS | 126516 | Biolegend |
| Pertussis Toxin | 100ng/ml | Glycerol | P2980 | Sigma |
| Rho kinase inhibitor | 1µM | Water | 555552 | Calbiochem |
| CCG215022 | 0.032-0.32 mM | DMSO | S6621 | Universal<br>biologicals |

Murine CTLs were used between d6-8 and were resuspended in 5% FCS complete RPMI on the top well. The reservoir plate was filled with 30% FCS complete RPMI containing 200nM Cxcl10 and 125nM Cxcl11 (Peprotech #250-16-250 and #250-29-250, respectively). Alternatively, the reservoir was seeded overnight with 6,000 MC38-OVA cells in 10% FCS DMEM/well. Just before the insert plate was added, the DMEM medium was aspirated and replaced by 30% FCS complete RPMI.

Human CTLs were used between d10-12 and were resuspended in Immunocult supplemented with 100 U/ml human IL-2. The reservoir plate was filled with the same media supplemented with 200nM CXCL10 and 125nM CXCL11 (Peprotech #300-12-100 and #300-46-100, respectively).

### RNA isolation

To validate the targeting of the Hh pathway by sonidegib in the tumour mouse cohorts, we collected a small piece of the gut, approximately 0.5 cm of the proximal duodenum. The tissue was washed briefly in PBS and flash frozen in Precellys lysis tubes (Precellys #432-3751) and subsequently homogenised in a Precellys Evolution Touch Homogeniser (3 cycles,15 sec, 6000 rpm, 4°C). RNA was extracted using the PureLink RNA Mini kit (Thermo #12183025) according to manufacturer’s instructions and RNA concentration was assessed by a Nanodrop spectrophotometer.

### qRT-PCR

Reactions for qRT-PCR were set up in 384 well plate format using the One-Step qRT-PCR Kit (Thermo Fisher SuperScript III Platinum #11732088) and Taqman probes (Thermo Fisher, Table 2), according to manufacturers’ instructions. Each sample was run in triplicate with *Tbp* serving as housekeeping gene. Reverse transcription thermal cycling was carried out on a QuantStudio 6 Flex Real-Time PCR System (Thermo Fisher).

**Table 2:** Probes used for qRT-PCR.

| Probe | Cat no. | Exon Boundary |
| --- | --- | --- |
| *CD3ε* | Mm00599684_g1 | 6-7 |
| *Tbp* | Mm00446973_m1 | 4-5 |
| *Smo* | Mm01162705_m1 | 2-3 |
| *Ptch1* | Mm00436026_m1 | 17-18 |
| *Gli1* | Mm00494654_m1 | 11-12 |
| *Ccl5* | Mm01302427_m1 | 1-2 |
| *Ccr1* | Mm01216147_m1 | 1-2 |
| *Ccr3* | Mm01216172_m1 | 1-2 |
| *Ccr5* | Mm01216171_m1 | 1-2 |
| *Cd160* | Mm00444461_m1 | 2-3 |
| *Cxcl9* | Mm00434946_m1 | 2-3 |
| *Cxcl10* | Mm00445235_m1 | 1-2 |
| *Cxcl11* | Mm00444662_m1 | 1-2 |
| *Cxcl12* | Mm00445553_m1 | 2-3 |
| *Cxcr4* | Mm01292123_m1 | 1-2 |
| *Icam1* | Mm00516023_m1 | 2-3 |
| *Ifn2/3* | Mm04204158_gH | 4-5 |
| *Gdf15* | Mm00442228_m1 | 1-2 |
| *Pdgfb* | Mm00440677_m1 | 4-5 |
| *PTCH1* | Hs00181117_m1 | 19-20 |
| *GLI1* | Hs01110766_m1 | 10-11 |
| *ACTINB* | Hs01060665_g1 | 2-3 |

Gene expression was calculated with the ΔCt method^46^. The cycle threshold (Ct) value from the gene of interest was subtracted from the housekeeping gene and transformed with a factor of 2^(−ΔCt) to give the fold expression relative to the housekeeping gene.

### Bulk RNA Seq

CD8 T cells were purified from spleens of *GzmBCreERT2/Rosa26eYFP/Smo* WT, HET and KO and stimulated for two days with plate-bound antibodies against CD3ε and CD28 in complete T cell media containing 300nM 4-OH-Tamoxifen (4-OHT) in presence of 100IU/ml IL-2. On d5, live CD8 eYFP positive and negative cells were FACS sorted, and cells were maintained in complete T cell media enriched with 50IU/ml IL-2. On d6, cells were harvested, and RNA was extracted using the miRNeasy mini kit (Qiagen #217004). RNA quality was assessed by 2100 Bioanalyzer Instrument (Agilent) using RNA 6000 Nano Kit (Agilent #5067-1511) and RNA concentrations were quantified using the Qubit RNA BR Assay kit (Thermo #10210). Libraries were generated using the TruSeq stranded mRNA kit (Illumina) and sequenced using single-read sequencing with the HiSeq4000 platform (Illumina). Reads were aligned to the mouse reference genome GRCm38132 using STAR v2.7.2b133 with default parameters and quality control was assessed using FASTQC v0.11.8 and Picard v2.9.5. Counts of reads against genes were generated using featureCounts from the Subread package v1.5.2134 and read counts were normalised and tested for differential gene expression using the DESeq2 package v1.5.2135.

### ConsensusTME

A small part of the tumour mass resected from the mouse cancer models was flash frozen in liquid nitrogen immediately after harvest. Samples were processed as previously described^28^. Gene expression was assessed by RNA-sequencing on a NovaSeq 6000 instrument. Reads were trimmed using Trimmomatic (v0.40)^47^ and alignment was performed using HiSat2 (v.2.2.1)^48^. ConsensusTME (v0.0.1.9000)^28^ was run with mouse gene sets derived from immunological genome project^49^ data to estimate immune cell abundance from mouse RNA-seq data.

### Western Blot

Cells were harvested at 4°C, washed twice in ice-cold PBS and lysed in ice-cold RIPA buffer (150mM NaCl, 50mM Tris pH 7.4, 1mM MgCl_2_, 2% NP40, 0.25% Na deoxycholate, 1mM DTT) with protease inhibitor cocktail (Pierce #88669). Protein concentration was determined with the Direct Detect spectrometer (Merck Millipore). Lysates were run on a NuPAGE 4-12% gradient Bis/Tris Acrylamide gel (Thermo Fisher #NP0335BOX). Western blotting was performed using wet transfer onto 0.45 µm nitrocellulose membrane (Thermo Fischer #LC2001) using the BioRad mini-transblot system. Membranes were blocked with 5% (w/v) non-fat dry milk (Marvel Original, Dried Skimmed Milk) in TBST and incubated with primary and secondary antibodies shown in Table 3. Membranes were incubated with stripping buffer (Restore PLUS Western Blot Stripping Buffer, Thermo Fisher #46430) between probing. Samples were visualised with SuperSignal West Pico Plus Chemiluminescent Substrate (Thermo Fisher #34580).

**Table 3:**
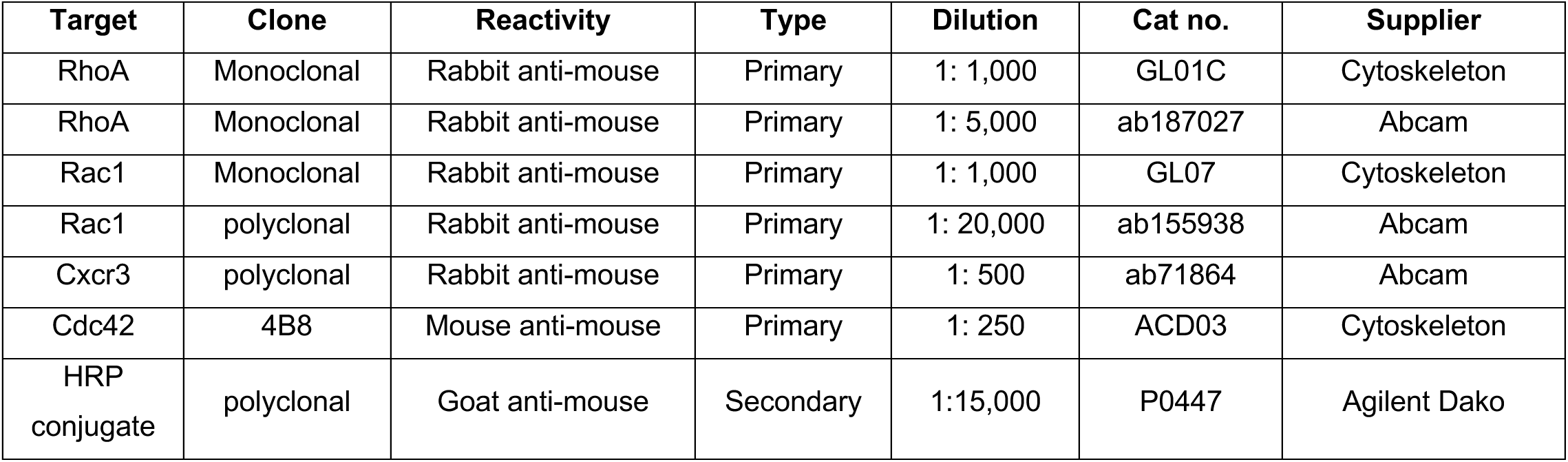
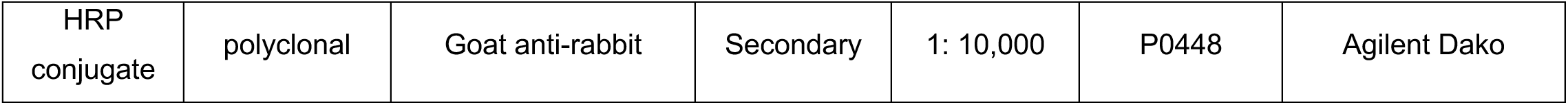
Antibodies used for western blot.

| Target | Clone | Reactivity | Type | Dilution | Cat no. | Supplier |
| --- | --- | --- | --- | --- | --- | --- |
| RhoA | Monoclonal | Rabbit anti-mouse | Primary | 1: 1,000 | GL01C | Cytoskeleton |
| RhoA | Monoclonal | Rabbit anti-mouse | Primary | 1: 5,000 | ab187027 | Abcam |
| Rac1 | Monoclonal | Rabbit anti-mouse | Primary | 1: 1,000 | GL07 | Cytoskeleton |
| Rac1 | polyclonal | Rabbit anti-mouse | Primary | 1: 20,000 | ab155938 | Abcam |
| Cxcr3 | polyclonal | Rabbit anti-mouse | Primary | 1: 500 | ab71864 | Abcam |
| Cdc42 | 4B8 | Mouse anti-mouse | Primary | 1: 250 | ACD03 | Cytoskeleton |
| HRP conjugate | polyclonal | Goat anti-mouse | Secondary | 1:15,000 | P0447 | Agilent Dako |
| HRP conjugate | polyclonal | Goat anti-rabbit | Secondary | 1: 10,000 | P0448 | Agilent Dako |

### Flow Cytometry

#### Surface staining

Cells were stained with fixable viability dye eFluor780 (eBioscience #65-0865-18), washed, and incubated with Fc block (1:100; Biolegend TruStain fcX #101320) and fluorophore-conjugated antibodies at the appropriate concentrations (Table 4) for 20min at 4°C. *Intracellular staining*: Following surface staining, cells were fixed with BD Cytofix/Cytoperm Plus Fixation Buffer (BD Biosciences #554715) as per manufacturer’s instructions and stained with fluorophore-conjugated antibodies for 30min at 4°C. Prior to analysis, cells were washed in permeabilization buffer. Flow cytometric analyses were conducted on a BD Fortessa or Symphony cell analyser, and data was analysed with FlowJo software (Tree Star Inc., v10.10).

**Table 4:**
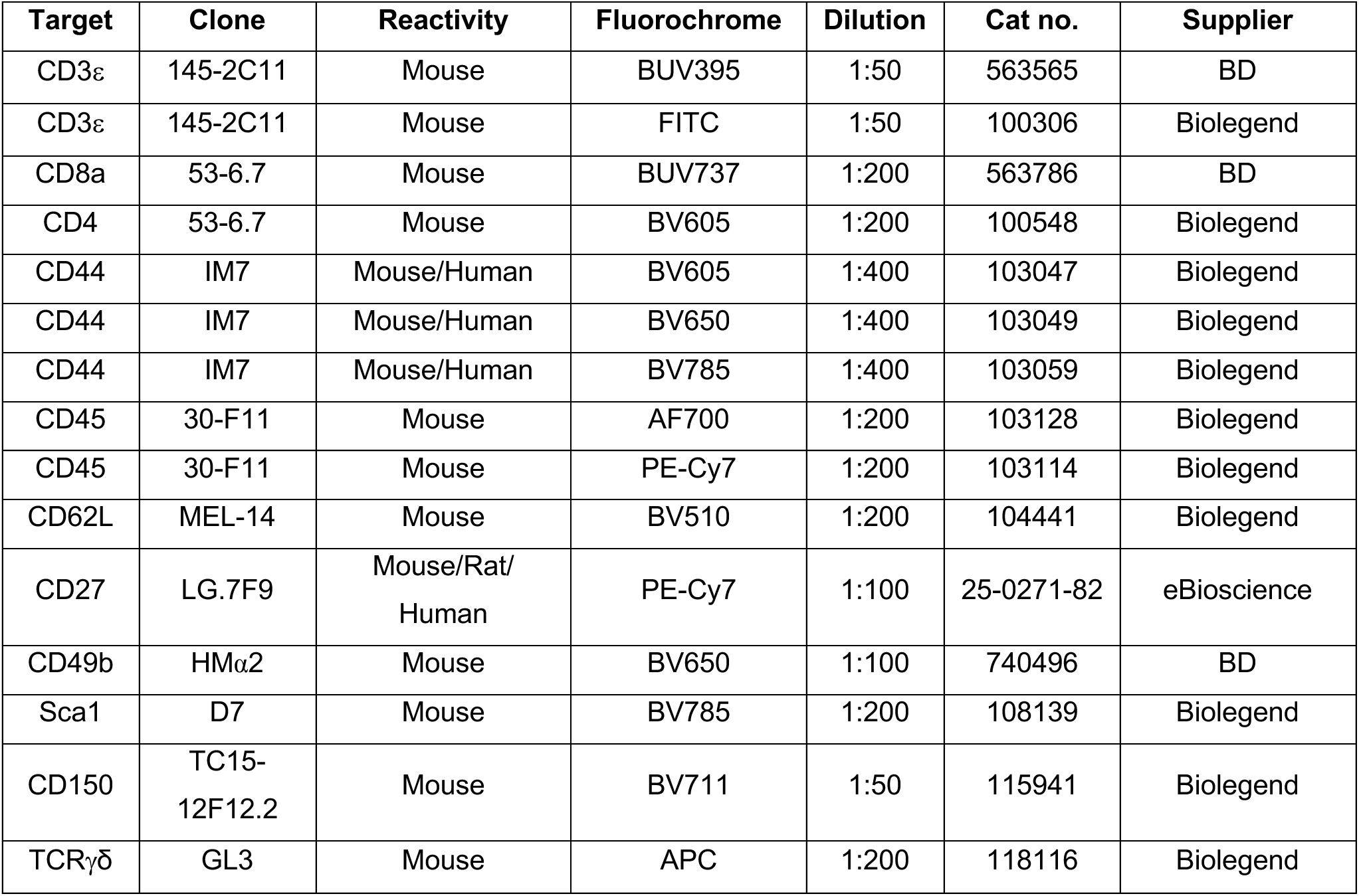
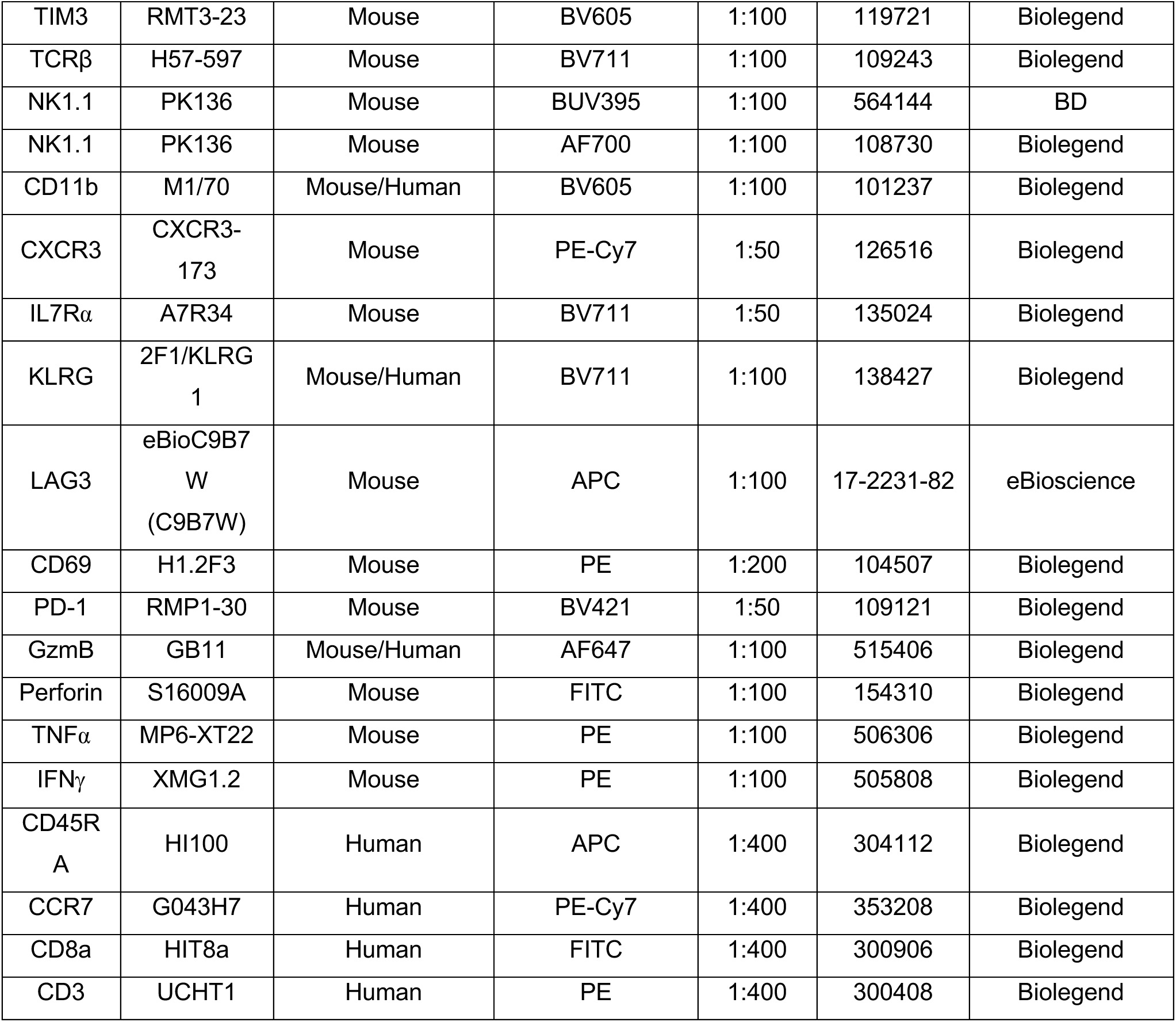
Antibodies used for flow cytometry.

| Target | Clone | Reactivity | Fluorochrome | Dilution | Cat no. | Supplier |
| --- | --- | --- | --- | --- | --- | --- |
| CD3ε | 145-2C11 | Mouse | BUV395 | 1:50 | 563565 | BD |
| CD3ε | 145-2C11 | Mouse | FITC | 1:50 | 100306 | Biolegend |
| CD8a | 53-6.7 | Mouse | BUV737 | 1:200 | 563786 | BD |
| CD4 | 53-6.7 | Mouse | BV605 | 1:200 | 100548 | Biolegend |
| CD44 | IM7 | Mouse/Human | BV605 | 1:400 | 103047 | Biolegend |
| CD44 | IM7 | Mouse/Human | BV650 | 1:400 | 103049 | Biolegend |
| CD44 | IM7 | Mouse/Human | BV785 | 1:400 | 103059 | Biolegend |
| CD45 | 30-F11 | Mouse | AF700 | 1:200 | 103128 | Biolegend |
| CD45 | 30-F11 | Mouse | PE-Cy7 | 1:200 | 103114 | Biolegend |
| CD62L | MEL-14 | Mouse | BV510 | 1:200 | 104441 | Biolegend |
| CD27 | LG.7F9 | Mouse/Rat/<br>Human | PE-Cy7 | 1:100 | 25-0271-82 | eBioscience |
| CD49b | HMα2 | Mouse | BV650 | 1:100 | 740496 | BD |
| Sca1 | D7 | Mouse | BV785 | 1:200 | 108139 | Biolegend |
| CD150 | TC15-12F12.2 | Mouse | BV711 | 1:50 | 115941 | Biolegend |
| TCR$\gamma\delta$ | GL3 | Mouse | APC | 1:200 | 118116 | Biolegend |
| TIM3 | RMT3-23 | Mouse | BV605 | 1:100 | 119721 | Biolegend |
| TCRβ | H57-597 | Mouse | BV711 | 1:100 | 109243 | Biolegend |
| NK1.1 | PK136 | Mouse | BUV395 | 1:100 | 564144 | BD |
| NK1.1 | PK136 | Mouse | AF700 | 1:100 | 108730 | Biolegend |
| CD11b | M1/70 | Mouse/Human | BV605 | 1:100 | 101237 | Biolegend |
| CXCR3 | CXCR3-173 | Mouse | PE-Cy7 | 1:50 | 126516 | Biolegend |
| IL7Rα | A7R34 | Mouse | BV711 | 1:50 | 135024 | Biolegend |
| KLRG | 2F1/KLRG1 | Mouse/Human | BV711 | 1:100 | 138427 | Biolegend |
| LAG3 | eBioC9B7W (C9B7W) | Mouse | APC | 1:100 | 17-2231-82 | eBioscience |
| CD69 | H1.2F3 | Mouse | PE | 1:200 | 104507 | Biolegend |
| PD-1 | RMP1-30 | Mouse | BV421 | 1:50 | 109121 | Biolegend |
| GzmB | GB11 | Mouse/Human | AF647 | 1:100 | 515406 | Biolegend |
| Perforin | S16009A | Mouse | FITC | 1:100 | 154310 | Biolegend |
| TNFα | MP6-XT22 | Mouse | PE | 1:100 | 506306 | Biolegend |
| IFNγ | XMG1.2 | Mouse | PE | 1:100 | 505808 | Biolegend |
| CD45RA | HI100 | Human | APC | 1:400 | 304112 | Biolegend |
| CCR7 | G043H7 | Human | PE-Cy7 | 1:400 | 353208 | Biolegend |
| CD8a | HIT8a | Human | FITC | 1:400 | 300906 | Biolegend |
| CD3 | UCHT1 | Human | PE | 1:400 | 300408 | Biolegend |

### Immunohistochemistry

Tissues were fixed in neutral buffered formalin (Sigma #HT5014) for 24hrs and then in 70% EtOH for 24hrs. Following fixation, the tissues were embedded in paraffin and sectioned at 3 μm in a Leica microtome. H&E and antibody staining was run on the Leica Bond-III platform according to CRUK-CI Histopathology facility protocols. The sodium citrate (Leica’s Epitope Retrieval Solution 1 #AR9961) and Tris EDTA (Leica’s Epitope Retrieval Solution 2 #AR9640) pre-treatments were performed at 100°C. The ancillary reagents are as follows: protein block from (Dako #X090930-2), anti-rat secondary (1:250, Bethyl Laboratories #A110-322A) and DAB Enhancer (Leica #AR9432) was applied to all antibodies. Antibodies and conditions are listed below:

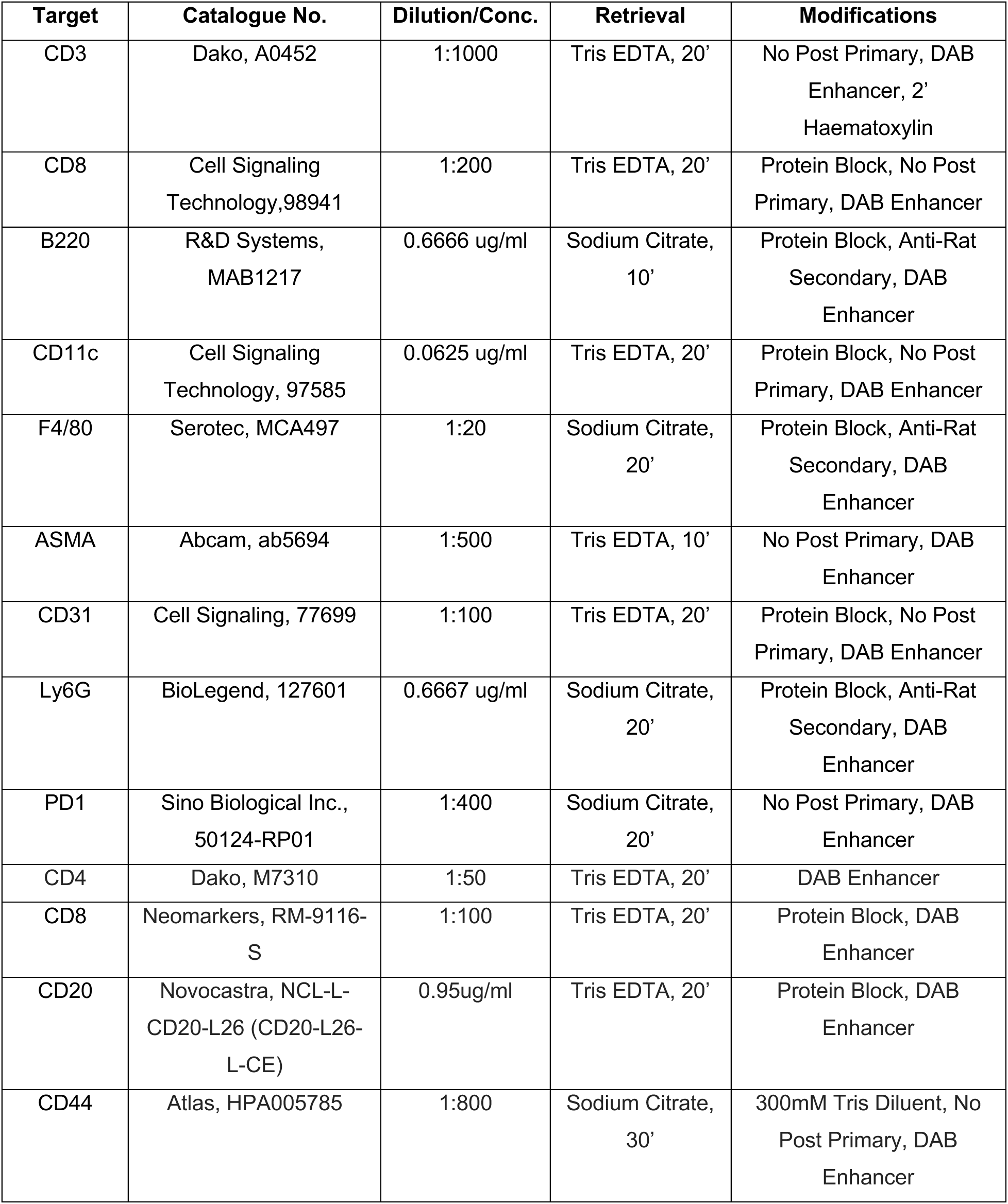
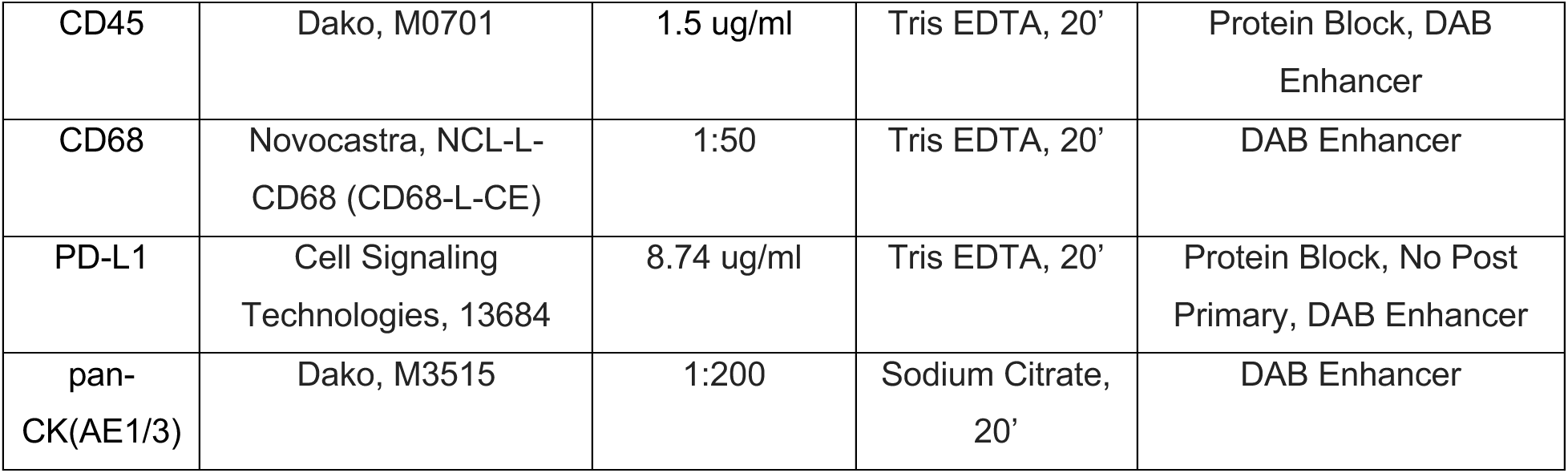

| Target | Catalogue No. | Dilution/Conc. | Retrieval | Modifications |
| --- | --- | --- | --- | --- |
| CD3 | Dako, A0452 | 1:1000 | Tris EDTA, 20' | No Post Primary, DAB Enhancer, 2' Haematoxylin |
| CD8 | Cell Signaling Technology, 98941 | 1:200 | Tris EDTA, 20' | Protein Block, No Post Primary, DAB Enhancer |
| B220 | R&D Systems, MAB1217 | 0.6666 ug/ml | Sodium Citrate, 10' | Protein Block, Anti-Rat Secondary, DAB Enhancer |
| CD11c | Cell Signaling Technology, 97585 | 0.0625 ug/ml | Tris EDTA, 20' | Protein Block, No Post Primary, DAB Enhancer |
| F4/80 | Serotec, MCA497 | 1:20 | Sodium Citrate, 20' | Protein Block, Anti-Rat Secondary, DAB Enhancer |
| ASMA | Abcam, ab5694 | 1:500 | Tris EDTA, 10' | No Post Primary, DAB Enhancer |
| CD31 | Cell Signaling, 77699 | 1:100 | Tris EDTA, 20' | Protein Block, No Post Primary, DAB Enhancer |
| Ly6G | BioLegend, 127601 | 0.6667 ug/ml | Sodium Citrate, 20' | Protein Block, Anti-Rat Secondary, DAB Enhancer |
| PD1 | Sino Biological Inc., 50124-RP01 | 1:400 | Sodium Citrate, 20' | No Post Primary, DAB Enhancer |
| CD4 | Dako, M7310 | 1:50 | Tris EDTA, 20' | DAB Enhancer |
| CD8 | Neomarkers, RM-9116-S | 1:100 | Tris EDTA, 20' | Protein Block, DAB Enhancer |
| CD20 | Novocastra, NCL-L-CD20-L26 (CD20-L26-L-CE) | 0.95ug/ml | Tris EDTA, 20' | Protein Block, DAB Enhancer |
| CD44 | Atlas, HPA005785 | 1:800 | Sodium Citrate, 30' | 300mM Tris Diluent, No Post Primary, DAB Enhancer |
| CD45 | Dako, M0701 | 1.5 ug/ml | Tris EDTA, 20' | Protein Block, DAB Enhancer |
| CD68 | Novocastra, NCL-L-CD68 (CD68-L-CE) | 1:50 | Tris EDTA, 20' | DAB Enhancer |
| PD-L1 | Cell Signaling Technologies, 13684 | 8.74 ug/ml | Tris EDTA, 20' | Protein Block, No Post Primary, DAB Enhancer |
| pan-CK(AE1/3) | Dako, M3515 | 1:200 | Sodium Citrate, 20' | DAB Enhancer |

Alternatively, human sections were processed as above and stained with a multiplex IF panel (Immuno8 FixVUE™ Panel including anti-human antibodies for: CD3, CD4, PD-1, CD8, PD-L1, CD11b, CD68, FOXP3 and CK/SOX10) according to manufacturer’s instructions.

### Immunohistochemistry analysis

To quantify CD8 infiltration into mouse tumours, serial tumour sections were stained with anti-mouse CD3 and CD8 as described above. Each tumour was subdivided in 100 μm-wide concentric rings and CD3+CD8+ double positive cells were enumerated per ring (HALO v3.6.4134.137).

### Live imaging

35mm Glass bottom dishes (MatTek #P35G-1.0-14-C) were coated with ICAM-1 (at 4°C, overnight at 1.25 μg/ml in PBS, BioTechne #796-IC-050). CTLs on days 6-8 after stimulation *in vitro* from *Smo* WT and KO or CTLs from *Rag1*KO OT-I mice were stained with CFSE (1:10,000 in PBS, Thermo #C34554) for 1h at 37°C. Where indicated, cells were pre-treated with cyclopamine overnight. CFSE-labelled CD8 T cells were washed and resuspended at 278,000 cells/ml in complete T cell media containing collagen (0.3 mg/ml PureCol Type I Bovine Collagen, Advanced Biomatrix #5005-100). Cell suspensions were added to the coated dishes and allowed to settle for 15 min in a humified chamber of an Andor Dragonfly 500 confocal spinning disc microscope (Oxford Instruments). Cells were imaged using the 40x oil immersion objective. Each picture is a composite of XY: 4×2 tiles, Z: 25 µm (2µm step size) and T: 20 min (20 sec interval).

### Image acquisition and analysis

Images were processed using Imaris software (Bitplane/Oxford Instruments). ‘Spots’ function was used to detect cells and the ‘tracking module’ function was used to map cell tracks. Mean speed, length and straightness of tracks were analysed for fully attached cells with tracks over 10 min long. 90-150 cell tracks were analysed per sample. The xy migration plots were based on cell tracks of speed >0.01 μm/sec using the spot function ‘plot all tracks with common origin’ from the Biological Imaging Development CoLab (BIDC) at UCSF Parnassus Heights.

### Clinical Trial Database

Results of published Hedgehog inhibitor clinical trials on ClinicalTrials.gov were compiled and ordered according to cancer type and colour-coded according to complete (CR) or partial (PR) objective response rates (ORRs) in patients (blue – positive effect; red – negative effect). Search terms used were ‘Sonidegib’, ‘Vismodegib’, ‘Arsenic Trioxide’ and ‘Itraconazole’. When both phases I and II are listed, the response rates and number of participants from phase II are reported. Number of patients exclusively includes number of patients whose response to treatment was evaluated (not number initially enrolled, if those differ from final number). Trials published up to April 2024 were included in the compiled data.

### Statistical analysis

For the statistical analysis of two groups, a two-sample unpaired t-test was performed. For analysis of multiple groups, one-way analysis of variance (ANOVA) with Tukey’s or multiple comparisons test was applied. For multiple testing, a two-way ANOVA with Sidak’s multiple comparisons test was applied. Family-wise significance and confidence level was set at P < 0.05. If not otherwise described, statistical examinations were carried out using GraphPad Prism (v.10.2.0). Additional information on the study design, the number of replicates and the statistical tests used is provided in the figure legends. Graphs and illustrations were created using GraphPad Prism and Affinity Designer (v.2.0.0).

## ACKNOWLEDGEMENTS

Special thanks and gratitude goes to Mike Mitchell, Michael Webb, Natalie Pettit, Gemma Cronshaw and Charlotte Gregg from the BRU for expert animal care and daily dosing; Andreas Bruckbauer, Fadwa Joud, Huw Naylor and Heather Zecchini from the Microscopy core for expert help and training; Jodi Miller and Cara Brodie from the Histology core for processing, staining and analysing tumour sections; Ashley Sawle from the Bioinformatics core for RNASeq analysis; and the flow cytometry, research instrumentation and compliance and biobanking cores for expert assistance; Lynn Cream, Doreen Milne and Sarah Loewenbein from the Cambridge Cancer Trials Centre for handling and archiving all the human samples and Nigel Burrows for his clinical insight; J. Hanna and James O. Jones for comments on the manuscript.

This work was supported by Cancer Research UK (MdlR, A22257), a Sir Henry Dale Fellowship jointly funded by the Wellcome Trust and the Royal Society (MdlR [107609/Z/15/Z]), a Wellcome Trust Discovery Award (MdlR [227432/Z/23/Z]) and the Addenbrooke’s Charitable Trust (KF and MdlR, Ref: 03/18 (v), Charity No.1170103).

## AUTHOR CONTRIBUTIONS

MdlR and CK conceived the project and designed the experiments. CK executed most experiments. LMOB performed all western blot and qRT-PCR analysis and supported all animal experiments. DS performed the review of the clinical trials using Hh inhibitors and performed live imaging. VC performed the bulk RNA-Seq experiment, OC performed Consensus TME analysis and SC-L and FB supported some experiments. SMcD is a consultant histopathologist who annotated the BCC sections. KF is a consultant oncologist and the clinical lead of the ALF trial, who co-designed the study, consented, and treated the BCC patients. CK and MdlR analyzed the results and wrote the manuscript. All authors commented on the manuscript.

