## Supplement for "Hedgehog signaling controls cytotoxic T cell migration in the tumour microenvironment"

### SUPPLEMENTAL MATERIAL

| Cancer Type | Drug | Study type | Patients | ORR | ORR Evaluation Criteria | Survival | NCT Number | Publications |
|---|---|---|---|---|---|---|---|---|
| Basal Cell Carcinoma (laBCC and mBCC) | Sonidegib | Randomised, double-blind, double-arm, phase II | 230 (94 - laBCC (66 receiving 200 mg, 128 receiving 800 mg) and 36 - mBCC (13 receiving 200 mg, 23 receiving 800 mg) | Final 42-month analysis:<br>56% (37/66) (200 mg) ; 46% (59/128) (800 mg) (laBCC) - 1.6% (800 mg) -5% (200 mg) CR<br>8% (1/13) (200 mg) -17% (4/23) (800 mg) (mBCC), 0% CR | RECIST | Median PFS: 22.1 (200mg) - 24.9 months (800mg) (laBCC); 13.1 (200mg) and 11.1 months (800 mg) months (mBCC).<br>Median follow-up: 38.2 months. | NCT01327053 | Migden et al, 2015 PMID: 25981810<br>Lear et al, 2018 PMID: 28846163<br>Dummer et al, 2020 PMID: 31545507 |
| | Vismodegib + Pembrolizumab | Non-randomised, open-label, double-arm, phase I/II | 16 (all mBCC) (9 - Pembrolizumab monotherapy; 7 - combination treatment) | At 18-week time point:<br>44% (4/9) - Pembrolizumab monotherapy arm<br>29% (2/7) - combination treatment arm | RECIST | 1-year PFS: 33% (3/9 ) - Pembrolizumab monotherapy arm<br>14% (1/7) - combination treatment arm | NCT02690948 | Chang et al, 2019<br>https://doi.org/10.1016/j.jaad.2018.08.017 |
| | Vismodegib | Open-label, single-arm, phase II | 1159 (1076 - laBCC; 83 - mBCC | 69% (745/1076) (laBCC), of which 33.9% are CR. 37% (31/83) (mBCC), of which 4.8% are CR. | RECIST | Median PFS: 20 months (laBCC) and 13 months (mBCC) | NCT01367665 | Basset-Seguin et al, 2015 PMID: 25981813<br>Basset-Seguin et al, 2017 PMID: 29073584<br>Bossi et al, 2020 PMID: 32374193 |
| | Vismodegib | Randomised, double-blind, regimen-controlled, phase II | 229 (116 - vismodegib intermittent schedule; 113 - vismodegib induction followed by intermittent schedule) | Overall ORR: 58% (133/229)<br>Intermittent schedule: 66% (76/116)<br>Induction followed by intermittent schedule: 50% (57/113)<br>(evaluated at data cut-off of 73 weeks after treatment start) | ≥50% reduction in total number of basal-cell carcinoma lesions from baseline | Not available | NCT01815840 | Jacobsen et al, 2017 PMID: 28325399<br>Dreno et al, 2017 PMID: 28188086 |
| | Vismodegib (neoadjuvant) | Open-label, non-comparative, phase II | 55 (all laBCC) | 80% (44/55) downstaging after Vismodegib.<br>ORR: 71% (39/55), of which 25.5% (14/55) are CR; 45.5% (25/55) PR. 29% (16/55) show SD. | RECIST | 22.7% (10/44) patients have ongoing responses 3 years after treatment start. 36% (16/44) had recurrence. | NCT02667574 | Bertrand et al, 2021 PMID: 33997740 |
| | Vismodegib | Non-randomised, open-label, phase I | 33 (15 - laBCC; 18 - mBCC) | 60% (2/15 - CR; 7/15 - PR) (laBCC) and 50% (9/18 - all PR) (mBCC) | RECIST | Median duration of response: 8.8 months | NCT00607724 | Von Hoff et al, 2009 PMID: 19726763 |
| | Vismodegib | Non-randomised, single-arm, two-cohort, phase II | 96 (63 - laBCC; 33-mBCC) | 60.3% (20/63 CR, 18/63 PR) laBCC<br>48.5% (16/33 PR) mBCC<br>Data cut-off: 39 months after completion of accrual. | RECIST | Median duration of response: laBCC - 26.2 months; mBCC - 14.8 months.<br>Median PFS: laBCC - 12.9 months; mBCC - 9.3 months.<br>Median OS: laBCC - not estimable; mBCC - 33.4 months. | NCT00833417 | Sekulic et al, 2012 PMID: 22670903<br>Chang et al, 2016 PMID: 27764798<br>Sekulic et al, 2017 PMID: 28511673 |
| | Itraconazole | Non-randomised, open-label, exploratory, double-arm, phase II | 21 (4 - Itraconazole 400 mg, 3 - Itraconazole 400 mg (prior Vismodegib); 4 - Itraconazole 200 mg; 10 - untreated). | 50% (4/8) PR in Itraconazole-only treated arm. The rest - SD<br>Tumour area decreased by 24%. | Visible decrease in tumour size, although a BCC still remains (partial response) | Not available | NCT01108094 | Kim et al, 2014<br>PMID: 24493717 |
| | Itraconazole (topical - 0.7%) | Open-label, double-arm phase II | 9 (6 - basal cell nevus syndrome; 3 - high-frequency BCCs) | No statistically significant difference in the percent change in tumor area between placebo- and topical itraconazole-treated tumours (evaluated at 4 and 12 weeks after treatment start). | change in BCC tumour area | Not available | NCT02735356 | Sohn et al, 2019<br>PMID: 31339515 |
| Medulloblastoma (recurrent/refractory) | Sonidegib | Open-label, single-arm, phase II | 39<br>(paediatric) | 5.1% (2/39) (paediatric) - all CR.<br>All responders are Hh-positive.<br>50% of patients with activated Hh pathway (5/10 - both adults and peadiatric) showed a response.<br>0% of patients (both adults and peadiatric) with a Hh-negative signature responded.<br>Hh pathway activation assessed by 5-gene Hh signature RT-PCR assay (Shou et al, 2015 - PMID: 25473003) | RECIST | Peadiatric CR duration: 21 months and the other was still in remission at the time the paper was written (49 months).<br>PFS: 7.6 months (paediatric) | NCT01125800 | Kieran et al, 2017<br>PMID: 28605510 |
| | | | 16<br>(adult) | 19% (3/16) (adult) - 2/16 are CR and 1/16 PR.<br>All responders are Hh positive.<br>Hh pathway activation assessed by 5-gene Hh signature RT-PCR assay (Shou et al, 2015 - PMID: 25473003) | RECIST | Adult CRs duration: 1.6 and 8.7 months; PR duration: 4.8 months.<br>PFS: 4.9 months (adult) | NCT01125800 | Kieran et al, 2017<br>PMID: 28605510 |
| | Vismodegib | Open-label, single-arm phase II | 12<br>(paediatric) | 8.3% (1/12) PR (SHH-MB) (response sustained for 8 weeks)<br>0% in non-SHH-MB<br>SHH subgroup classification based on IHC. | CR - disappearance of all target lesions; PR - >= 50% reduction in tumour size. | Median PFS:<br>1.41 months<br>Duration of objective response: 2.8 months | NCT01239316 | Robinson et al, 2015<br>PMID: 26169613 |
| | Vismodegib | Open-label, single-arm phase II | 31<br>(adult) | Overall ORR: 9.7% (3/31) (responses sustained for 8 weeks)<br>0% (0/8)- SHH pathway inactivated<br>0% (0/3)- SHH pathway activation unknown<br>15% (3/20) -SHH pathway activated<br>SHH subgroup classification based on IHC. | CR - disappearance of all target lesions; PR - >= 50% reduction in tumour size. | PFS: 1.64 months - SHH pathway inactivated<br>1.48 months - SHH pathway activation unknown<br>2.76 months - SHH pathway activated<br>Median duration of objective response: 4.6 months | NCT00939484 | Robinson et al, 2015<br>PMID: 26169613 |
| | Vismodegib + Temozolomide | Randomised, open-label, three-arm, phase I/II | 24 (adult) (10-TMZ + Vismodegib; 5 - TMZ alone; 9 - Vismodegib alone (previously treated with TMZ) | 40%(4/10) PR - TMZ + Vismodegib<br>20% (1/5) PR - TMZ alone<br>22.2% (2/9) PR - Vismodegib alone (previous treatment with TMZ)<br>All MB with documented SHH-pathway activation - based on IHC. | RECIST | 6-month PFS is not significantly increased by the addition of Vismodegib to TMZ compared to TMZ-alone - led to study termination.<br>Median PFS: 4.8 months - TMZ + Vismodegib<br>3.8 months - TMZ alone; 1.9 months - Vismodegib with TMZ previously. | NCT01601184 | Frappaz et al, 2021<br>PMID: 33825892 |
| Triple-negative advanced Breast | Sonidegib + Docetaxel | Open-label, single arm phase I | 10 | 10% (1/10) (800 mg) - CR<br>20% (2/10) - SD | RECIST | Time to Progression: 203+ days for CR; 155 and 188 days for SD. | NCT02027376 | Ruiz-Borrego et al, 2018<br>PMID: 29948356 |
| Myelofibrosis | Saridegib | Open-label, single-arm, phase II | 12 | All patients discontinued treatment by 7.5 months.<br>75% (9/12) -no response<br>17% (2/12) - developed acute leukemia.<br>8% (1/12) - disease progression | International Working Group for Myelofibrosis Research and Treatment (IWG-MRT) criteria | Not available | NCT01371617 | Sasaki et al, 2015<br>PMID: 25641433 |
| | Vismodegib + ruxolitinib | Open-label, single-arm Phase I | 8 | After 48 weeks of treatment:<br>12.5% (1/8) showed clinical improvement<br>75% (6/8) - SD<br>12.5% (1/8) relapsed (previously- PR)<br>No evidence of increased efficacy with Vismodegib compared to Ruxolitinib alone. | IWG-MRT criteria | Not available | NCT02593760 | Couban et al, 2018<br>PMID: 30249277 |
| | Sonidegib+ ruxolitinib | Open-label, single-arm, phase I/II, | 27 (phase II) | Sonidegib + ruxolitinib:<br>Week 48: 29.6%<br>Ruxolitinib only (result from phase III COMFORT trials):<br>Week 48: 28%<br>No further development of combination as modest overall benefit compared with historical ruxolitinib monotherapy.<br>(JAK inhibitor-naïve myelofibrosis patients) | IWG-MRT criteria: ≥35% reduction in spleen volume (MRI/CT) | Not available | NCT01787552 | Gupta et al, 2020<br>PMID: 32634234 |
| Extensive stage small cell lung cancer | Arsenic Trioxide (ATO) | Open-label, single-arm, phase II | 17 | 0% CR/PR.<br>12% (2/17) SD<br>88% (15/17) PD<br>(relapsed SCLC) | RECIST | PFS: 6.3 weeks<br>OS:4.5 months | NCT01470248 | Owonikoko et al, 2016<br>PMID: 27142472 |
| | Sonidegib+ Etoposide + cisplatin (EP) chemotherapy | Open-label, single-arm phase I | 14 | 79% (11/14) PR<br>21% (3/14) SD | RECIST | Median PFS: 5.5 months; Median OS: 19.7 months These are consistent with expected outcomes of EP-alone.<br>One patient with SOX2 amplification remains progression-free on maintenance Sonidegib after 27 months (unusual for EP-alone). | NCT01579929 | Pietanza et al, 2016<br>PMID: 27565909 |
| | Vismodegib/cixutumab + Etoposide + cisplatin (EP) chemotherapy | Randomised, open-label, three-arm, phase II | 152 | 48% (23 /48) - EP<br>56% (29/52) - EP + vismodegib<br>50% (26/52)- EP + cixutumab<br>Not statistically significant. | RECIST | Median PFS:<br>4.7 months - EP<br>4.4 months - EP + vismodegib<br>4.6 mo nths - EP + cixutumab<br>Not statistically significant. | NCT00887159 | Prakash Belani et al, 2013<br>https://doi.org/10.1200/jco.2013.31.15_suppl.7508 |

| Cancer Type | Drug | Study type | Patients | ORR | ORR Evaluation Criteria | Survival | NCT Number | Publications |
|---|---|---|---|---|---|---|---|---|
| Non-small cell lung cancer | Itraconazole + gemcitabine chemotherapy combination | Randomised, open-label, prospective, double-arm phase II | 60 (30 per arm) | 90% - Itraconazole + chemotherapy<br>66.7% - Chemotherapy<br>(significant difference)<br>(chemotherapy-naïve metastatic NSCLC) | RECIST | Mean 1-year PFS: 5.4 months - Chemotherapy<br>6.1 months - Itraconazole + chemotherapy<br>No significant difference in 1-year OS. | NCT03664115 | Mohamed et al, 2021<br>PMID: 33559053 |
| Prostate cancer | Vismodegib | Single-arm, early phase I | 9 | No patient achieved a PSA reduction or a measurable tumor response.<br>100% PSA Increase.<br>(castration-resistant metastatic prostate cancer) | RECIST | At the time of data cutoff, all patients have progressed and six patients have died.<br>Median PFS: 1.87 months.<br>Median OS: 7.04 months | NCT02115828 | Maughan et al, 2016<br>PMID: 27826729 |
| | Itraconazole | Randomised, noncomparative, phase II | 46 (29 -600 mg;17 - 200 mg) | 0% (200 mg) and 14.3% (600 mg)<br>Itraconazole - favourable effects on CTC count.<br>(chemotherapy-naïve metastatic castration-resistant prostate cancer) | PSA response: ≥50% decline in PSA levels maintained for ≥ 4 weeks. | Median PSA PFS (PPFS) at 24 weeks: 11.8% (low-dose arm); 48% (high-dose arm)<br>Median PFS: 11.9 weeks (low-dose arm) and 35.9 weeks (high-dose arm).<br>(PSA progression: ≥25% increase in PSA from nadir). | NCT00887458 | Antonarakis et al, 2013<br>PMID: 23340005 |
| | Sonidegib (neoadjuvant prior to prostatectomy) | Randomised, open-label, prospective, double-arm phase I | 14 (7 per arm) | Sonidegib: median PSA increase of + 0.4 ng/mL<br>Control (observation-only): median PSA decrease of -1 ng/mL.<br>(high-risk localised prostate cancer) | PSA levels | At the time of data cutoff, with a median follow-up of 181.5 days, disease progression (PSA ≥ 0.2 ng/mL) occurred in:<br>Sonidegib: 57% (4/7)<br>Control: 29% (2/7) | NCT02111187 | Ross et al, 2017<br>PMID: 29262631 |
| Ovarian cancer | Itraconazole + chemotherapy (docetaxel + carboplatin + gemcitabine) | Retrospective Study | 55 patients (19 - chemotherapy and intraconazole; 36 - chemotherapy alone) | 32% (6/19)- Chemotherapy + Itraconazole:<br>11%(4/36)- Chemotherapy alone<br>(refractory ovarian cancer) | RECIST | Median PFS: 53 days - Chemotherapy<br>103 days - Chemotherapy + itraconazole<br>Median OS:139 days - Chemotherapy<br>642 days - Chemotherapy + itraconazole | Not available | Tsubamoto et al, 2014<br>PMID: 24778064 |
| | Vismodegib (maintenance therapy) | Randomised, double-blind, double-arm, phase II | 104 (52 in each arm) | Not available<br>(ovarian cancer in second or third CR) | Not available. | Median PFS: 7.5 months - Vismodegib<br>5.8 months - Placebo<br>The intended increase in PFS was not achieved. | NCT00739661 | Kaye et al, 2012<br>PMID: 23032746 |
| Advanced Chondrosarcoma | Vismodegib | Open-label, single-arm, phase II | 39 | 0% ORR<br>SD at 6 months after inclusion: 25.6% (10/39)<br>PD: 74.4% (29/39)<br>Predefined 6-month CBR (objective) of 40% was not met. | RECIST<br>(CBR - CR, PR, SD) | Median PFS: 3.5 months<br>OS: 12.4 months.<br>Median follow-up: 13.9 months. | NCT01267955 | Italiano et al, 2013<br>PMID: 24170610 |
| Glioblastoma multiforme (recurrent) | Vismodegib (neoadjuvant) | Randomised, open-label, double-arm phase II | 40 (20 per arm) | 0% CR or PR.<br>PD: 76.5 (13/17) - pre-surgery vismodegib; 68.8% (11/16) - no pre-surgery vismodegib.<br>SD: 20% (4/20) - pre-surgery vismodegib; 25% (5/20) - no pre-surgery vismodegib: | Macdonald Radiographic Response Criteria | 6-month PFS: Pre-surgery vismodegib: 0% (0/20)<br>No pre-surgery vismodegib: 5% (1/20)<br>OS: Pre-surgery vismodegib: 7.8 months<br>No pre-surgery vismodegib: 7.6 months | NCT00980343 | Sloan et al, 2014 |
| Advanced Sarcoma | Vismodegib + Notch signalling pathway Inhibitor (RO4929097) | Randomised open-label phase I/II (study prematurely terminated due to RO4929097 discontinuation) | 67 (34 - Notch inhibitor alone; 33 - Notch inhibitor + vismodegib) | no ORR; SD: 50% (17/34) - Notch inhibitor only<br>51.5% (17/33) - Notch inhibitor + Vismodegib | RECIST | Median PFS (not significant):<br>8.9 weeks - Notch inhibitor only<br>12.0 weeks - Notch inhibitor + Vismodegib<br>The median follow-up for survivors was 42.8 months. | NCT01154452 | Gounder et al, 2012<br>https://doi.org/10.1158/1078-0432.CCR-21-3874 |
| Refractory B cell Lymphoma and CLL | Vismodegib | Open-label Phase II | 31 (12 - diffuse large B-cell lymphoma; 6 - 'indolent' lymphoma; 10 - primary central nervous system lymphoma; 3 - CLL) | 3% (1/31 ('indolent' follicular lymphoma) - PR (highest expression of GLI by PCR).<br>ORR for lymphomas alone: 3.6% (1/28).<br>All patients discontinued treatment prematurely due to disease progression (90.3%). | According to Cheson 1999 for DLBCL and iNHL, IPCG response criteria for PCNSL and IWCLL response criteria for CLL. | Median PFS: 1.7 months.<br>PR in 1 patient lasted for 4.8 months. | NCT01944943 | Houot et al, 2016<br>https://doi.org/10.1093/annonc/mdw138 |
| Acute Leukemia | Glasdegib and Cytarabine Chemotherapy (low dose) (LDAC) | Randomised, open-label, double-arm phase II | 116 (78 combination arm; 38 LDAC alone) | CR: Glasdegib + LDAC: 19.2% (15/78);<br>LDAC: 2.6% (1/38)<br>(newly diagnosed or secondary AML, unsuitable for intensive chemotherapy) | International Working Group response criteria (IWG) + WHO Guidelines | Median OS (statistically significant):<br>Glasdegib + LDAC: 8.3 months<br>LDAC: 4.3 months<br>Median duration of CR: 379.5 days with Glasdegib+LDAC; 91 days with LDAC alone.<br>Longest follow-up since randomisation: 36 months. | NCT01546038 | Cortes et al, 2019<br>Heuser et al, 2021 PMID: 33740113 |
| | Sonidegib | Randomised, openl-label, double-arm phase II | 70 (35 in each dosing arm (400 mg and 800 mg)) | Complete remission with incomplete blood count recovery: 1.4% (1/35 in 400 mg arm; 0/35 in 800 mg arm)<br>(relapsed/refractory acute leukemia) | International Working Group response criteria (IWG) | Not available | NCT01826214 | Not available. |
| Pancreatic cancer | Vismodegib + Gemcitabine | Pilot single-arm phase II | 23 | 21.7% (5/23) PR; 60.9% (14/23) SD; 39.1% (9/23) PD.<br>No statistically significant decrease of CSCs and no additional clinical benefit compared to Gemcitabine alone.<br>(metastatic pancreatic ductal adenocarcinoma) | RECIST | Median PFS: 2.8 months Median OS: 5.3 months<br>Vismodegib and Gemcitabine did not improve median PFS or OS compared with historical data for Gemcitabine alone. | NCT01195415 | Kim et al, 2014<br>PMID: 25278454 |
| | Vismodegib + Sirolimus | Open-label, single-arm phase I | 22 | 0% - CR/PR<br>(metastatic pancreatic adenocarcinoma) | RECIST | Not available | NCT01537107 | Carr et al, 2020<br>https://doi.org/10.1016/j.pan.2020.06.015 |
| | Vismodegib + Gemcitabine | Randomised, double-blind, double-arm phase II | 106 - 53 in each arm | Gemcitabine + Vismodegib: 8% (4/53) (all PR); 51% (27/53) SD.<br>Gemcitabine + Placebo: 13% (7/53) (1 CR; 6 PR); 38% (20/53) SD.<br>No statistically significant difference.<br>(recurrent or metastatic pancreatic cancer) | RECIST | OS:Gemcitabine + Vismodegib: 6.9 months<br>Gemcitabine + Placebo: 6.1 months<br>No statistically significant difference. | NCT01064622 | Catenacci et al, 2015<br>PMID: 26527777 |
| | Vismodegib + Gemcitabine + Nab-Paclitaxel | Open-label, single-arm phase II | 67 | 40% (27/67) had a response (1 CR; 38.8% (26/67) - PR.<br>No improvement compared to historical chemotherapy alone.<br>No significant change in CSCs.<br>(untreated metastatic pancreatic adenocarcinoma) | RECIST | Median PFS: 5.42 months<br>Median OS: 9.79 months. | NCT01088815 | De Jesus-Acosta et al, 2019<br>PMID: 31857726 |
| | Sonidegib + Gemcitabine | Open-label, dingle-arm phase I | 18 | 11% (2/18) - ORR; 61% (11/18) PD<br>(locally advanced or metastatic pancreatic cancer) | Not available | Median PFS - 4.9 months<br>No improvement in PFS compared to current standard. | NCT01487785 | Macarulla et al., 2016<br>https://doi.org/10.1200/jco.2016.34.4_suppl.371 |
| | Sonidegib + Gemcitabine and Nab-paclitaxel | Open-label, single-arm, phase II | 19 | 10% (2/19) PR; 53%(10/19) SD; 37% (7/19) PD<br>(metastatic pancreatic cancer - prior chemotherapy) | RECIST | Median OS 6 months. | NCT02358161 | Pijnappel et al, 2019 - https://doi.org/10.1093/annonc/mdz247.022 |
| | Saridegib + FOLFIRINOX | Open-label, single-arm phase I | 15 | 66.7% (9/15) - unconfirmed OR<br>As the trial was accruing, early results available from Infinity Pharmaceuticals phase II trial (see below). | RECIST | Median PFS - 8.4 months. | NCT01383538 | Ko et al, 2016<br>PMID: 26390428 |
| | Saridegib (+ Gemcitabine | Randomised, double-arm phase II | Not reported | More rapid rate of disease progression in Gemcitabine-treated vs placebo-arm - study voluntarily halted. | Not reported | Shorter median survival time in Gemcitabine-treated vs placebo arm. | n/a (Trial sponsored by Infinity Pharmaceuticals ) | Infinity Pharmaceuticals website: http://phx.corporate-ir.net/phoenix.zhtml?c=121941&p=irol-newsArticle_print&ID=1653550&highlight= |
| Advanced Gastro-esophageal cancer | Vismodegib + FOLFOX Chemotherapy | Randomised, double-blind, double-arm phase II | 124 (64 - placebo arm: 60 - Vismodegib arm) | 37% (22/60) - FOLFOX and Vismodegib<br>44% (28/64) - FOLFOX and Placebo | RECIST | OS: 12.12 months - FOLFOX and Vismodegib<br>15.4 months - FOLFOX and Placebo: | NCT00982592 | Cohen et al, 2013<br>https://doi.org/10.1200/jco.2013.31.4_suppl.67 |
| Metastatic colorectal cancer | Vismodegib + FOLFOX/FOLFIRI Chemotherapy + Bevacizumab | Randomised, double-blind, double-arm phase II | 199 (101 - placebo arm; 98 - Vismodegib arm) | 46% (45/98) - Chemotherapy + Bevacizumab + Vismodegib<br>51% (52/101) - Chemotherapy + Bevacizumab + Placebo | RECIST | Median PFS: 9.3 months - Chemotherapy + Bevacizumab + Vismodegib<br>10.1 months - Chemotherapy + Bevacizumab + Placebo<br>Median follow-up: 12.6 months | NCT00636610 | Berlin et al, 2013 PMID: 23082002 |

**Supplemental 1: Detailed overview of clinical trials involving Hedgehog inhibitors in cancer patients.**

Results of published Hedgehog inhibitor clinical trials on ClinicalTrials.gov were compiled according to cancer type. When applicable, a more detailed description of the cancer (and any prior treatments) is specified in brackets at the end of column ‘ORR’. Search terms used were ‘Sonidegib’, ‘Vismodegib’, ‘Arsenic Trioxide’ and ‘Itraconazole’. When both phases I and II are listed, the response rates and number of participants from phase II are reported. Number of patients exclusively includes number of patients whose response to treatment was evaluated (not number initially enrolled, if those differ from final number). N/A: not available/assessed, CR: complete response, PR: partial response, PFS: progression-free survival, OS: overall survival, SD: stable disease, PD: progressive disease. *Link will be provided as a searchable database.*

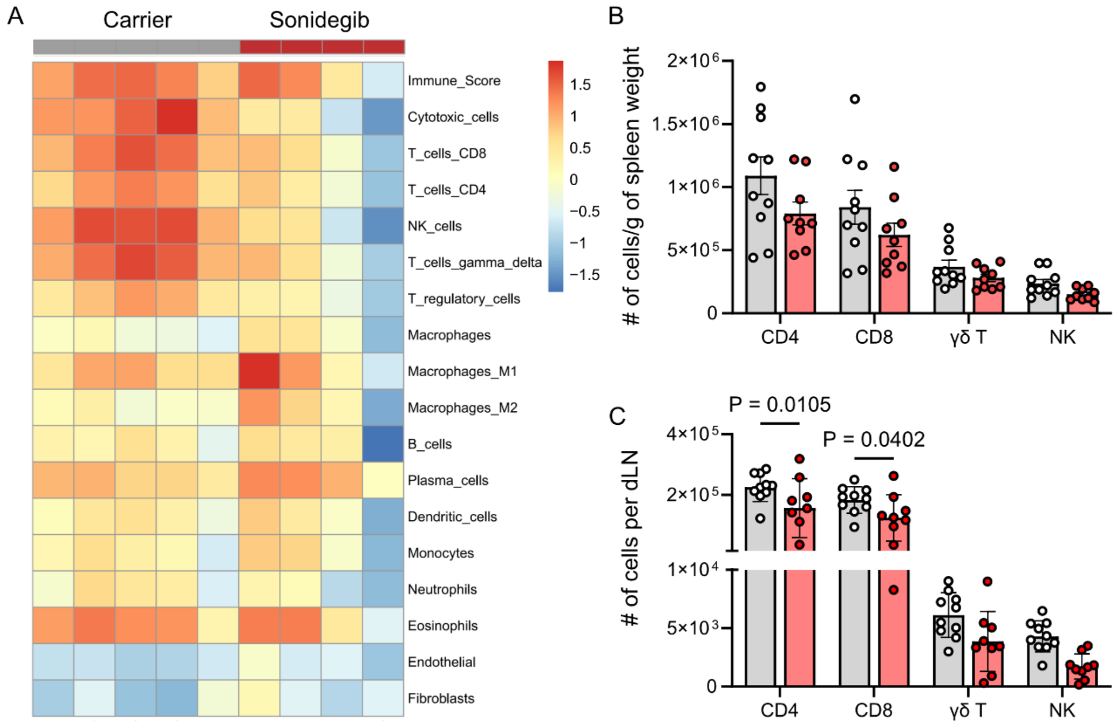


**Supplemental 2: Sonidegib treatment diminishes the lymphoid compartment in the tumour.**

**(A)** Bulk RNA-Seq data of MC38 tumours from mice treated with either carrier or sonidegib analysed by Consensus-TME. n=5 for carrier-treated tumours, n=4 for sonidegib-treated tumours, one independent experiment.
**(B)** Numbers of CD4+, CD8+ and gammadelta T cells as well as NK cells in the spleens of MC38 tumour-bearing mice measured by flow cytometry and shown per gram of spleen weight. n=10 for carrier-treated mice, n=9 for sonidegib-treated mice, two independent experiments, two-way ANOVA, mean ± SEM.
**(C)** Numbers of CD4+, CD8+ and gamma delta T cells as well as NK cells in the tumour-draining lymph nodes of MC38 tumour-bearing mice measured by flow cytometry. n=10 for carrier-treated mice, n=9 for sonidegib-treated mice, two independent experiments, two-way ANOVA, mean ± SEM.

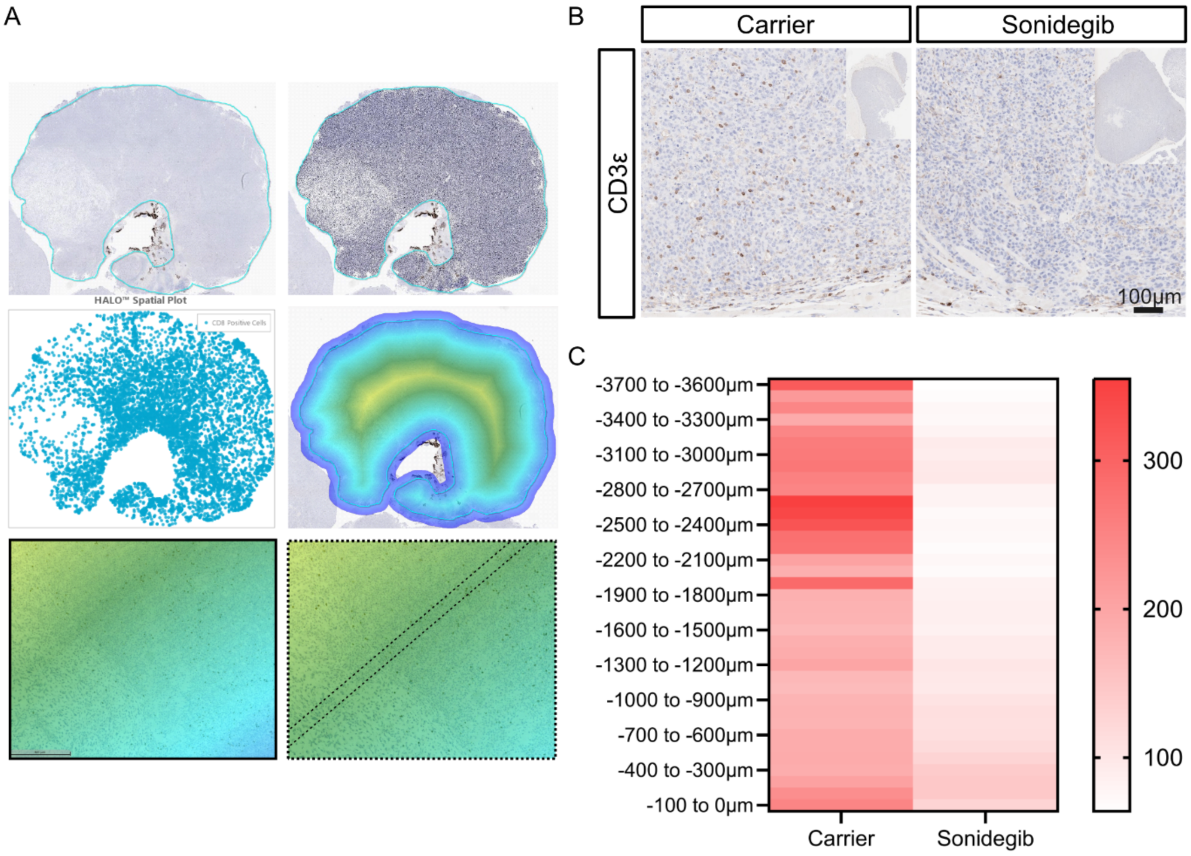


**Supplemental 3: Infiltration analysis in MC38 tumours treated with sonidegib.**

**(A)** Workflow of T cell infiltration analysis using Halo software. First mask was applied to outline the tumour margins and a second mask to indicate CD3/CD8 double positive cells. Subsequently, the tumour was subdivided in 100 µm concentric zones starting from the tumour border towards the centre of the tumour. Such analysis was performed in **Fig. 3C** and **Suppl. Fig. 6C, 8G, 8N**.

**(B)** Representative paraffin sections of MC38 tumours from mice treated either with carrier control or sonidegib and stained with anti-mouse CD3ε antibodies. The whole tumour is shown in top right insert panels.

**(C)** Numbers of CD3+ cells/ $mm^2$ in 100µm-wide zones from the tumour surface (-100 to 0 µm) to the tumour centre (-3700 to -3600 µm) as quantified by HALO analysis (workflow shown in (**A**). Mean is shown. n=10 for carrier-treated and n=8 for sonidegib-treated mice, two independent experiments.

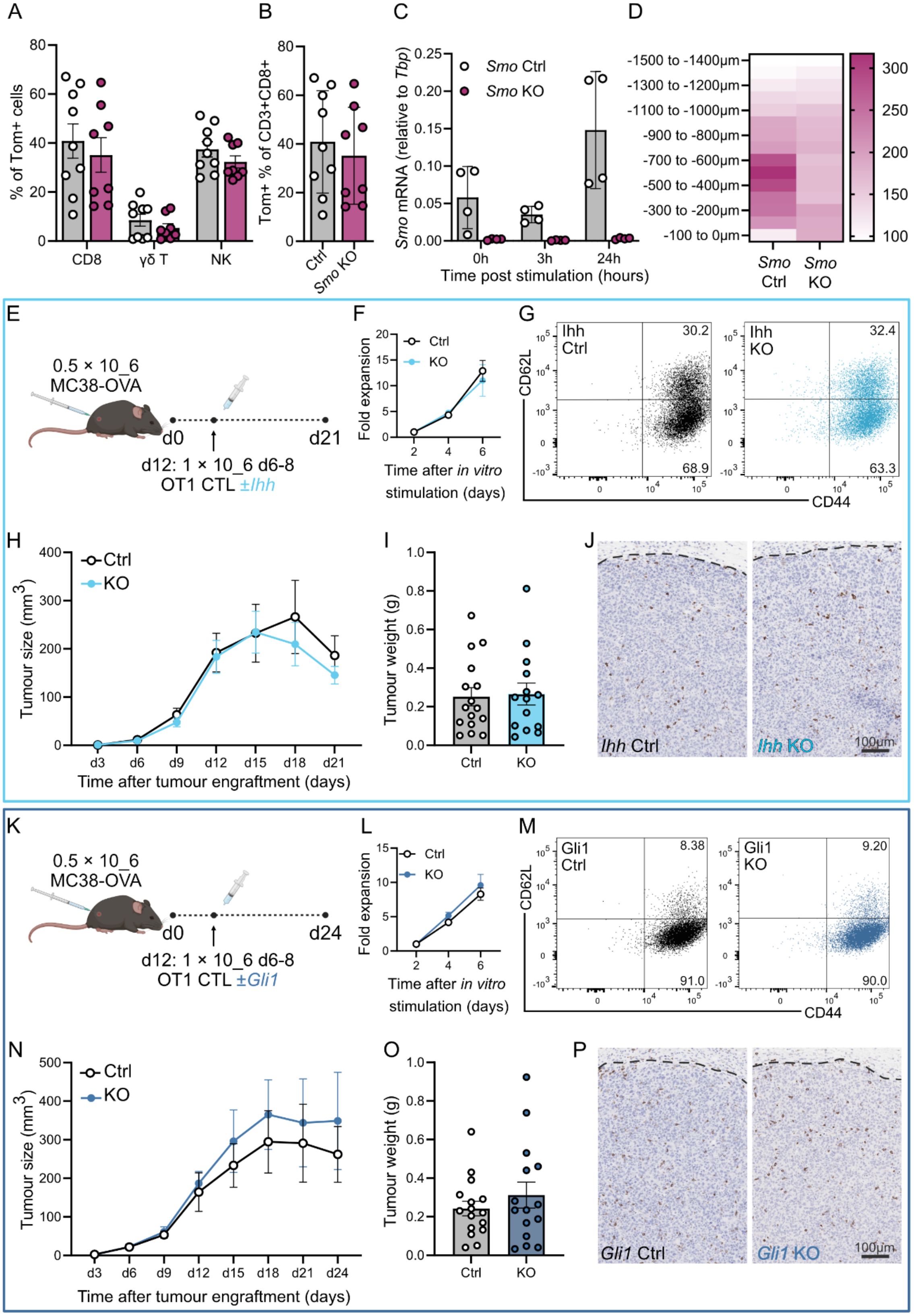
A
% of Tom+ cells
CD8
γδ T
NK
B
Tom+ % of CD3+CD8+
Ctrl
Smo KO
C
Smo mRNA (relative to Tbp)
Smo Ctrl
Smo KO
0h
3h
24h
Time post stimulation (hours)
D
-1500 to -1400µm
-1300 to -1200µm
-1100 to -1000µm
-900 to -800µm
-700 to -600µm
-500 to -400µm
-300 to -200µm
-100 to 0µm
Smo Ctrl
Smo KO
E
0.5 × 10_6 MC38-OVA
d0
d21
d12: 1 × 10_6 d6-8 OT1 CTL ±Ihh
F
Fold expansion
Ctrl
KO
Time after in vitro stimulation (days)
G
Ihh Ctrl
Ihh KO
CD62L
CD44
30.2
68.9
32.4
63.3
H
Tumour size (mm3)
Ctrl
KO
Time after tumour engraftment (days)
I
Tumour weight (g)
Ctrl
KO
J
Ihh Ctrl
Ihh KO
100µm
K
0.5 × 10_6 MC38-OVA
d0
d24
d12: 1 × 10_6 d6-8 OT1 CTL ±Gli1
L
Fold expansion
Ctrl
KO
Time after in vitro stimulation (days)
M
Gli1 Ctrl
Gli1 KO
CD62L
CD44
8.38
91.0
9.20
90.0
N
Tumour size (mm3)
Ctrl
KO
Time after tumour engraftment (days)
O
Tumour weight (g)
Ctrl
KO
P
Gli1 Ctrl
Gli1 KO
100µm

50

**Supplemental 4: Genetic loss of *Smo* but not *Ihh* or *Gli1* in cytotoxic CD8 cells does affect tumour infiltration and growth.**

**(A)** *GzmB-ERT2Cre/ROSAtdTom/Smo*$^{fl/fl}$ (KO) or *Smo*$^{fl/+}$ (Ctrl) animals were injected with 0.5 × $10^6$ MC38 cells subcutaneously on d0. Mice were treated with tamoxifen via intraperitoneal injections (75mg/kg) on -d1, d1, d3 and d5. Percentage of immune subsets in the tumour microenvironment in which *Smo* has been excised after tamoxifen administration. Four independent experiments, n=9 for *Smo*$^{fl/+}$ (Ctrl), n=8 for *Smo*$^{fl/fl}$ (KO).
**(B)** Intratumoural CD8s from (**A**) were assessed for tdTom expression by flow cytometry at endpoint. Four independent experiments, n=9 for *Smo*$^{fl/+}$ (Ctrl), n=8 for *Smo*$^{fl/fl}$ (KO).
**(C)** Expression of *Smo* mRNA in CD8 T cells isolated from *GzmB-ERT2Cre/ROSAtdTom/Smo*$^{fl/fl}$ (KO) or *Smo*$^{fl/+}$ (Ctrl) animals after *ex vivo* expansion and restimulation on d10 at indicated timepoints. OHT was added to the cultures for the first 5 days and *Tbp* was used as a housekeeping gene. Similar results were obtained when *CD3e* was used as a reference gene. n=4 for *Smo*$^{fl/+}$ (Ctrl), n=4 for *Smo*$^{fl/fl}$ (KO), two independent experiments.
**(D)** Quantification of T cell infiltration shown in **Fig.4J** by HALO analysis. Two independent experiments, n=4 for *Smo*$^{fl/+}$ (Ctrl), n=10 for *Smo*$^{fl/fl}$ (KO), mean is shown.
**(E)** Experimental Design. *Rag2*KO animals were subcutaneously injected with 0.5 × $10^6$ MC38-OVA cells on d0. On d12, mice were stratified into two equal groups according to tumour size. In parallel, single cell suspensions from spleens and lymph nodes of *dLck-Cre/Ihh*$^{fl/fl}$ (KO) or *Ihh*$^{fl/+}$ (Ctrl) OTI mice were stimulated *in vitro* with OVA peptide for 48hrs and subsequently cultured for 6-8 days.
**(F)** Fold expansion of CD8 cells during *in vitro* culture.
**(G)** Representative graphs of CD62L/CD44 expression of CTLs on d7 used for the adoptive transfers (**H**, **I** of this figure).
**(H)** Tumour dimensions were determined by caliper measurements. n=16 for *Ihh*$^{fl/+}$ (Ctrl), n=14 for *Ihh*$^{fl/fl}$ (KO), two independent experiments, ordinary two-way ANOVA with Geisser-Greenhouse correction, mean ± SEM.
**(I)** Tumour weight at endpoint (d21) from (**H**), unpaired Mann-Whitney test, mean ± SEM.
**(J)** Representative paraffin sections from MC38-OVA tumours shown in (**H**, **I**) and stained with anti-mouse CD8$\alpha$ antibodies. Dotted line indicates tumour margins.
**(K)** Experimental Design. *Rag2*KO animals were subcutaneously injected with 0.5 × $10^6$ MC38-OVA cells on d0. On d12, mice were stratified into two equal groups according to tumour size. In parallel, single cell suspensions from spleens and lymph nodes of *Gli1*$^{eGFP/eGFP}$ (KO) or *Gli1*$^{+/+}$ (Ctrl) OTI mice were stimulated *in vitro* with OVA peptide for 48hrs and subsequently cultured for 6-8 days.
**(L)** Fold expansion of CD8 cells during *in vitro* culture.
**(M)** Representative graphs of CD62L/CD44 expression of CTLs on d7 used for the adoptive transfers (**N**, **O** of this figure).
**(N)** Tumour dimensions were determined by caliper measurements. n=12 for *Gli1*$^{+/+}$ (Ctrl), n=11 for *Gli1*$^{eGFP/eGFP}$ (KO), three independent experiments, ordinary two-way ANOVA with Geisser-Greenhouse correction, mean ± SEM.
**(O)** Tumour weight at endpoint (d21) from (**M**), unpaired Mann-Whitney test, mean ± SEM.
**(P)** Representative paraffin sections from MC38-OVA tumours shown in (**N**, **O**) and stained with anti-mouse CD8$\alpha$ antibodies. Dotted line indicates tumour margins.

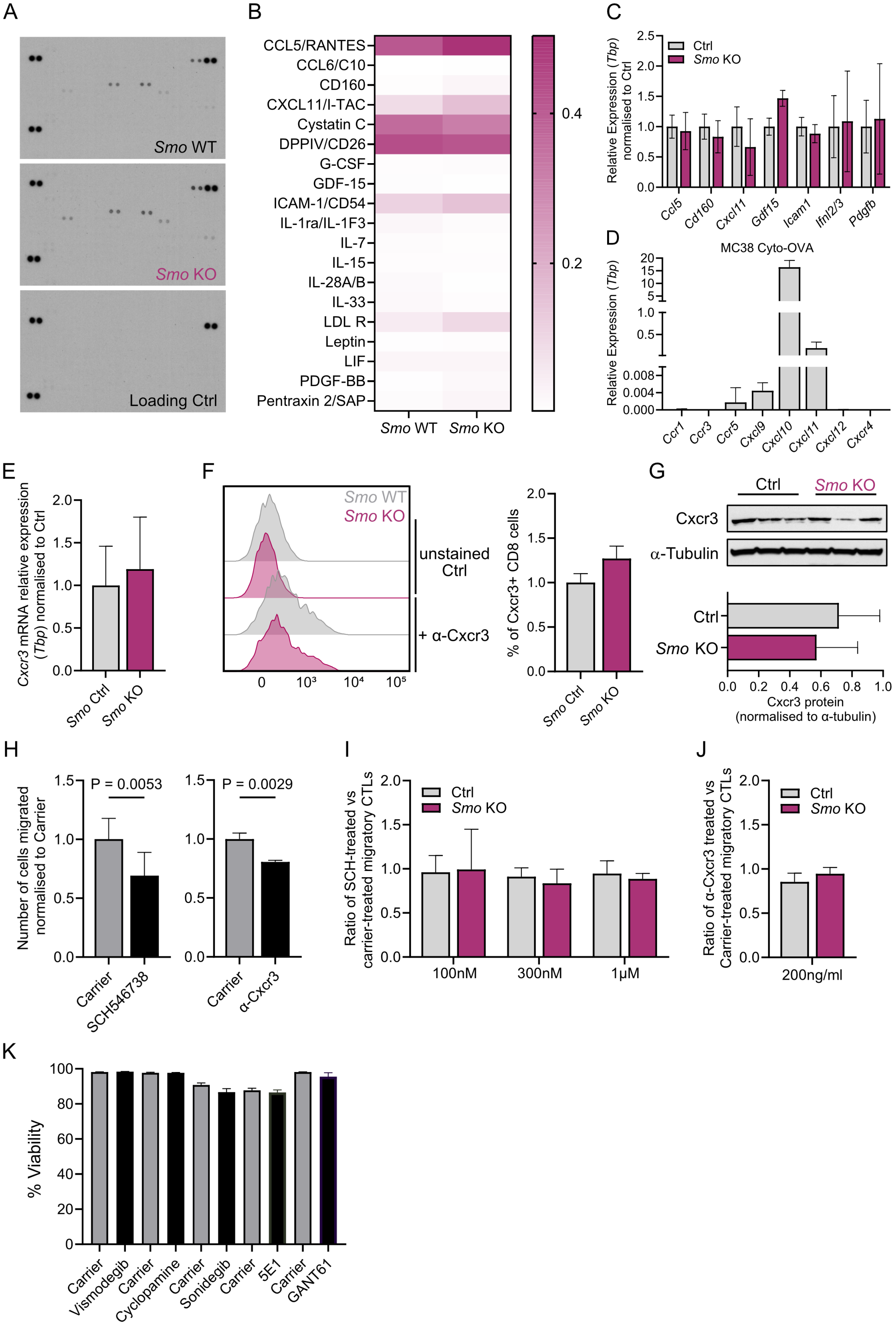
A
Smo WT
Smo KO
Loading Ctrl
B
CCL5/RANTES
CCL6/C10
CD160
CXCL11/I-TAC
Cystatin C
DPPIV/CD26
G-CSF
GDF-15
ICAM-1/CD54
IL-1ra/IL-1F3
IL-7
IL-15
IL-28A/B
IL-33
LDL R
Leptin
LIF
PDGF-BB
Pentraxin 2/SAP
Smo WT
Smo KO
0.4
0.2
C
Relative Expression (Tbp) normalised to Ctrl
Ctrl
Smo KO
Ccl5
Cd160
Cxcl11
Gdf15
Icam1
Ifnl2/3
Pdgfb
D
MC38 Cyto-OVA
Relative Expression (Tbp)
Ccr1
Ccr3
Ccr5
Cxcl9
Cxcl10
Cxcl11
Cxcl12
Cxcr4
E
Cxcr3 mRNA relative expression (Tbp) normalised to Ctrl
Smo Ctrl
Smo KO
F
Smo WT
Smo KO
unstained Ctrl
+ α-Cxcr3
% of Cxcr3+ CD8 cells
Smo Ctrl
Smo KO
G
Ctrl
Smo KO
Cxcr3
α-Tubulin
Cxcr3 protein (normalised to α-tubulin)
H
Number of cells migrated normalised to Carrier
P = 0.0053
P = 0.0029
Carrier
SCH546738
α-Cxcr3
I
Ratio of SCH-treated vs carrier-treated migratory CTLs
100nM
300nM
1μM
J
Ratio of α-Cxcr3 treated vs Carrier-treated migratory CTLs
200ng/ml
K
% Viability
Carrier
Vismodegib
Cyclopamine
Sonidegib
5E1
GANT61

**Supplemental 5: Genetic loss of *Smo* in cytotoxic CD8 cells does not affect general cytokine production profiles or response to tumour chemoattractants via the Cxcr3-Cxcl9/10/11 axis.**

CTLs were generated from *GzmB-ERT2Cre/ROSAtdTom* $Smo^{fl/fl}$ (KO) and $Smo^{+/+}$ or $Smo^{f/+}$ (Ctrl) mice and used for downstream assays shown in (**A-C, E-G, I-K**) between d7/8 of *ex vivo* culture.

**(A)** Profiler XL array for cytokines expressed by $Smo^{fl/fl}$ (KO) or $Smo^{+/+}$ (Ctrl) CTLs on d8 lysed cell pellets.

**(B)** Quantification of signal intensity from (**A**). The mean of two technical repeats is shown in a single-gradient heatmap. One independent experiment, n=1 for *Smo* WT and n=1 for *Smo* KO.

**(C)** mRNA expression levels of highly expressed targets from the XL profiler array (**A, B**) assessed by qRT-PCR with *Tbp* as a housekeeping gene. n=4 for *Smo* Ctrl and n=3 for *Smo* KO, one independent experiment, mean ± SD.

**(D)** mRNA levels of key chemokines expressed by MC38-OVA cells. Four pellets were included and run in triplicates and *Tbp* was used as a housekeeping gene. One independent experiment, mean ± SD.

**(E)** mRNA expression levels of *Cxcr3* assessed by qRT-PCR, n=4 for *Smo* Ctrl and n=3 for *Smo* KO, one independent experiment, unpaired t-test showed no significant difference, P=0.7724, mean ± SD.

**(F)** Left: Representative flow cytometry histograms for Cxcr3 staining. Right: Percentage of Cxcr3+ cells out of CD3+CD8+ cells on d8. n=7 for *Smo* Ctrl and n=7 for *Smo* KO, two independent experiments, unpaired t-test showed no significant difference, P=0.1426, mean ± SD.

**(G)** Cxcr3 protein expression analysed by western blot with α-Tubulin used as a loading control. One independent experiment, n=3 for *Smo* Ctrl and n=3 for *Smo* KO. For the quantification panel (bottom), mean + SD is shown, P=0.5408. Two additional independent experiments have been performed with n=3 for $Smo^{fl/+}$ or $Smo^{+/+}$ (Ctrl) and n=3 for $Smo^{fl/fl}$, also showing no significant differences between Ctrl and *Smo* KO (data not shown).

**(H)** Transwell assays with Cxcl10 and Cxcl11-supplemented T cell media at the bottom well and d7/8 CTLs in drug-containing media in the insert wells. After 6hrs, cells from the bottom well were collected for flow cytometric analysis. SCH546738 (Cxcr3 antagonist) 100nM, n=7, two independent experiments. α-Cxcr3 200ng/ml, n=11, four independent experiments. For each drug paired t-tests were performed, and mean ± SEM is shown.

**(I)** Transwell assays as in (**H**). Ratios of SCH546738-treated compared to carrier-treated CTLs is shown for each concentration. For 100nM, n=4 for *Smo* Ctrl and n=3 for *Smo* KO from two independent experiments. For 300nm and 1µM, n=4 for *Smo* Ctrl and n=4 for *Smo* KO, one independent experiment, mean ± SD.

**(J)** Transwell assays as in (**H**). Ratios of α-Cxcr3-treated (200ng/ml) compared to carrier-treated CTLs is shown. Two independent experiments, n=7 for *Smo* Ctrl and n=7 for *Smo* KO, mean ± SD.

**(K)** Viability of murine CD8+ cells after treatment with the drugs used for transwell assays in **Fig. 5B, C.**

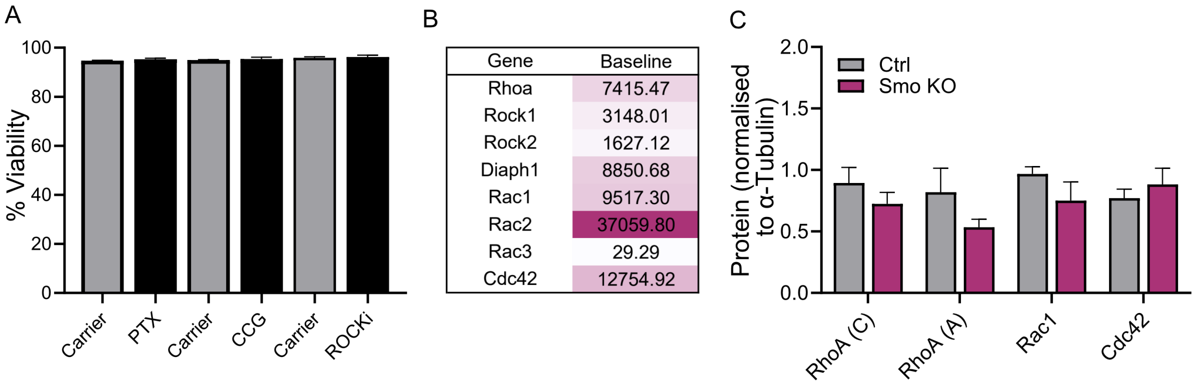


**Supplemental 6: Steady state levels of RhoA, Rac1, and Cdc42 are unchanged upon genetic loss of *Smo*.**

**(A)** Viability of human CD8+ T cells after treatment with the drugs used for transwell assays in **Fig. 6B, C, F**. Two independent experiments, n=6 mice, mean + SEM is shown. **(B)** mRNA expression of small GTPases determined by bulk RNASeq performed on murine CTLs on day 6 post stimulation, n=7 mice. **(C)** Quantification of protein levels of SDS-PAGE gels shown in **Fig. 6D**. Values are normalised to loading control, α-Tubulin. Two separate antibodies were tested for RhoA, one form Cytoskeleton (C) and one from Abcam (A). One independent experiment, n=3 for $Smo^{fl/+}$ (Ctrl), n=3 for $Smo^{fl/fl}$ (KO), mean + SD is shown. Two additional independent experiments have been performed with n=3-4 for $Smo^{fl/+}$ or $Smo^{+/+}$ (Ctrl) and n=3-4 for $Smo^{fl/fl}$, also showing no significant differences between Ctrl and *Smo* KO (data not shown).

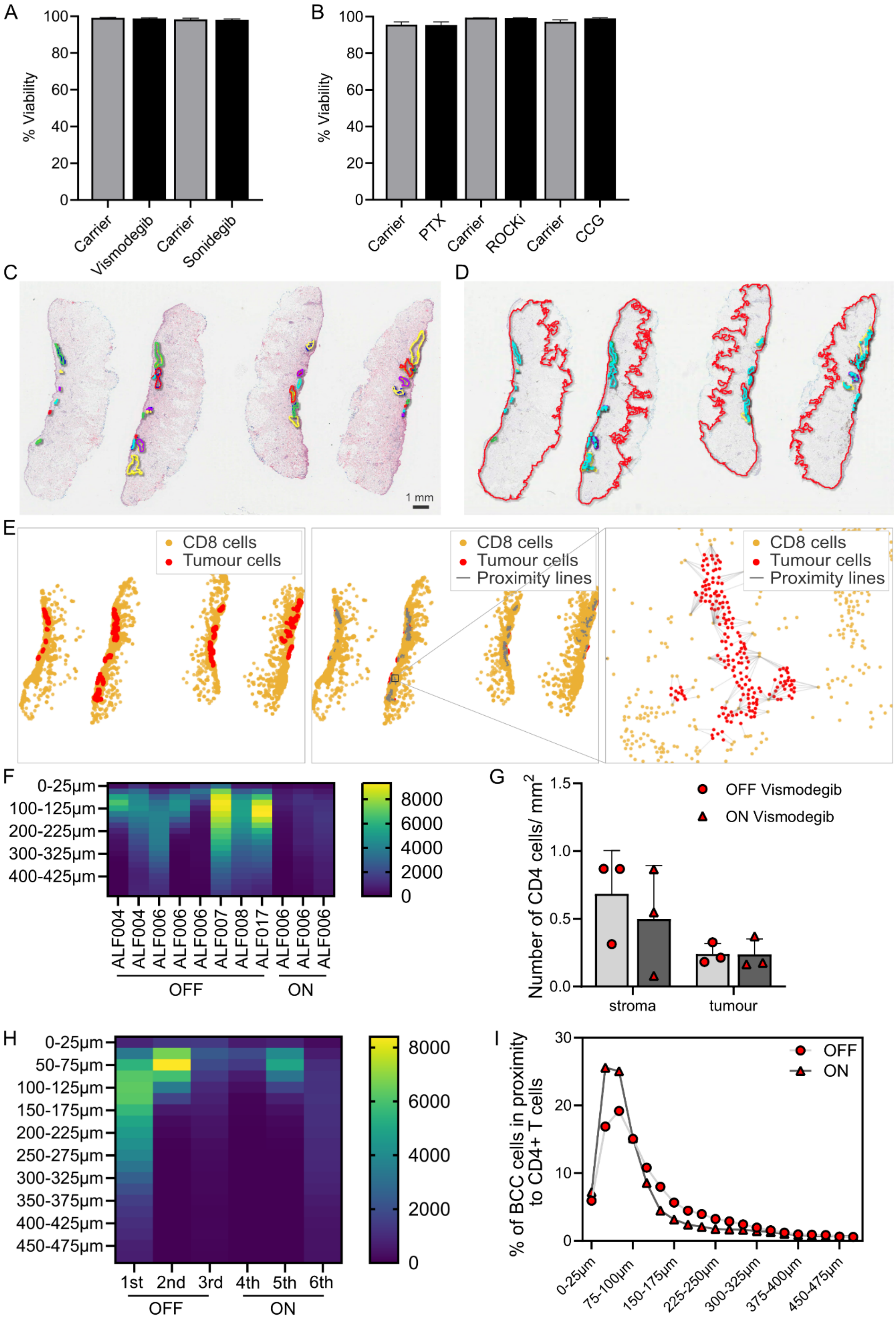
A
% Viability
Carrier
Vismodegib
Carrier
Sonidegib
B
% Viability
Carrier
PTX
Carrier
ROCKi
Carrier
CCG
C
1 mm
D
E
CD8 cells
Tumour cells
Proximity lines
F
0-25µm
100-125µm
200-225µm
300-325µm
400-425µm
ALF004
ALF004
ALF006
ALF006
ALF006
ALF007
ALF008
ALF017
ALF006
ALF006
ALF006
OFF
ON
G
OFF Vismodegib
ON Vismodegib
Number of CD4 cells/ mm2
stroma
tumour
H
0-25µm
50-75µm
100-125µm
150-175µm
200-225µm
250-275µm
300-325µm
350-375µm
400-425µm
450-475µm
1st
2nd
3rd
4th
5th
6th
OFF
ON
I
% of BCC cells in proximity to CD4+ T cells
OFF
ON
0-25µm
75-100µm
150-175µm
225-250µm
300-325µm
375-400µm
450-475µm

**Supplemental 7: Image analysis of BCC biopsies shows diminished T cell infiltration upon vismodegib treatment.**

**(A)** Viability of human CD8+ T cells after treatment with the drugs used for transwell assays in **Fig. 7A**.

**(B)** Viability of human CD8+ T cells after treatment with the drugs used for transwell assays in **Fig. 7B**. Only the highest concentration (0.32mM) tried for CCG215022 is shown.

**(C-E)** Workflow of imaging analysis for human BCC sections.

**(C)** Annotation of tumour margins in H&E-stained BCC biopsies performed by a clinical histopathologist.

**(D)** Consecutive section of (**C**) stained with antibodies against human CD8a with classifiers for tumour and stroma regions shown.

**(E)** Mask of CD8 T cells in yellow and tumour cells in red with every dot representing one cell (left), same mask with proximity lines drawn in grey between each BCC cell and its nearest CD8 T cell neighbour (middle) with magnification of box region shown (right).

**(F)** Proximity analysis of each BCC cell to its closest neighbouring CD8+ T cell. Heatmap indicates the number of BCC tumour cells within the indicated distance brackets. All biopsies we were able to obtain through our clinical "ALF" study are shown including the ones shown in **Fig. 7F**. n=3 BCCs *"on"* and n=8 BCCs *"off"* vismodegib treatment from 5 patients in total.

**(G)** Numbers of CD4+ T cells found in the stroma (defined as within 500 µm outwards of the tumour margin) or the tumour (defined as within the tumour margins) per $mm^2$ of stroma and tumour areas, respectively. n=3 *"on"* and n=3 *"off"* treatment, mean ± SD is shown.

**(H)** Proximity analysis of each BCC cell to its closest neighbouring CD4+ T cell. Heatmap indicates the number of BCC tumour cells within the indicated distance brackets. Separate resected tumour specimens of the same patient (ALF006) spanning 11 years while the patient was *"on"* or *"off"* vismodegib treatment were analysed. n=3 *"on"* and n=3 *"off"* treatment.

**(I)** Percentage of total BCC cells shown with regards to their closest neighbouring CD4+ T cell (analysis workflow shown in **Supp. Fig. 7C-E**) adjusted for tumour load. Means of n=3 BCCs *"on"* and n=3 BCCs *"off"* treatment are shown.

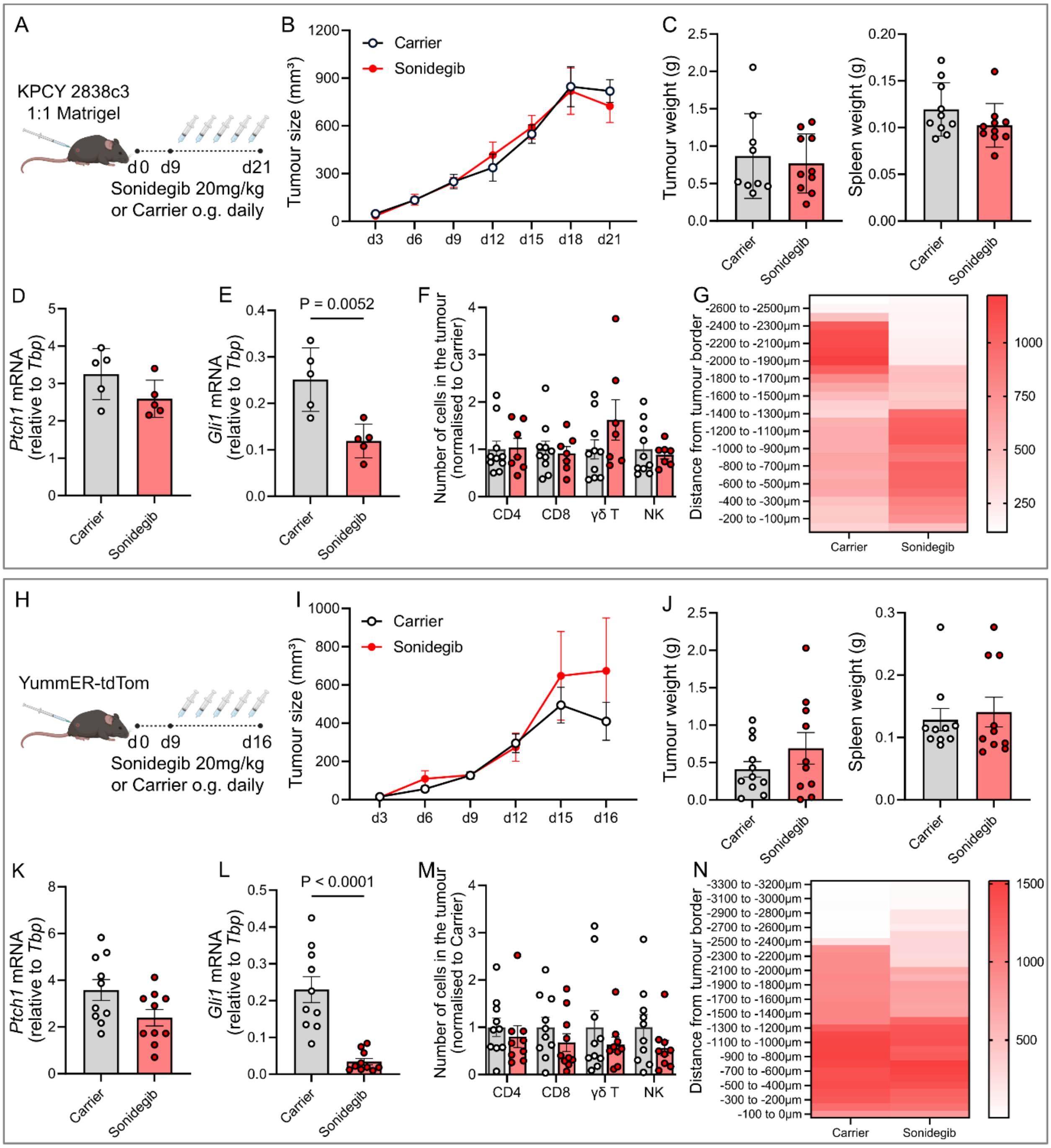


**Supplemental 8: Sonidegib treatment fails to reduce the tumour burden in murine models of pancreatic cancer or melanoma.**
**(A)** Experimental design. C57BL/6J wildtype animals were subcutaneously injected with $1 \times 10^6$ KPCY-2838c3 pancreatic cancer cells on d0 at a 1:1 ratio with matrigel. On d9, mice were stratified into two equal groups according to tumour size. Between d9 and d21, mice were treated daily with 20mg/kg sonidegib or carrier control by oral gavage.

**(B)** Tumour dimensions as established by caliper measurements. Two independent experiments, n=10 for carrier-treated mice, n=10 for sonidegib-treated mice, ordinary two-way ANOVA with Geisser-Greenhouse correction, mean ± SEM.
**(C)** Tumour and spleen weight at endpoint (d21). Two independent experiments, n=10 for carrier-treated mice, n=10 for sonidegib treated mice, unpaired Mann-Whitney test, mean ± SEM.
**(D, E)** mRNA levels of *Ptch1* and *Gli1* in the small intestine. One independent experiment, n=5 for for carrier-treated mice, n=5 for sonidegib treated mice, unpaired Mann-Whitney test, mean ± SEM.
**(F)** Enumeration of immune subsets in the tumour by flow cytometry at endpoint (d21), normalised to the carrier controls. Two independent experiments, n=10 for carrier-treated mice, n=7 for sonidegib treated mice, multiple unpaired t-tests, mean ± SEM.
**(G)** Mean of number of CD8+ cells/ $mm^2$ in 100μm-wide zones from the tumour surface (-100 to 0 μm) to the tumour centre (-2600 to -2500 μm) as quantified by HALO software (**Suppl. Fig. 3A**). Two independent experiments, n=10 for carrier treatment mice, n=10 for sonidegib-treated mice. Mean is shown in a single gradient heatmap.
**(H)** Experimental design. C57BL/6J wildtype animals were subcutaneously injected with $1 \times 10^6$ YummER-tdTom melanoma cells on d0. On d9, mice were stratified into two equal groups according to tumour size. Between d9 and d16, mice were treated daily with 20mg/kg sonidegib or carrier control by oral gavage.
**(I)** Tumour dimensions as established by caliper measurements. Two independent experiments, n=12 for carrier-treated mice, n=13 for Sonidegib treated mice, ordinary two-way ANOVA with Geisser-Greenhouse correction, mean ± SEM.
**(J)** Tumour and spleen weight at endpoint (d16). Two independent experiments, n=12 for carrier-treated mice, n=10 for sonidegib treated mice, unpaired Mann-Whitney test, mean ± SEM.
**(K, L)** mRNA levels of *Ptch1* and *Gli1* in the small intestine. Two independent experiments, n=10 for carrier-treated mice, n=10 for sonidegib treated mice, unpaired Mann-Whitney test, mean ± SEM.
**(M)** Flow cytometry enumeration of immune subsets in the tumour at endpoint (d21), normalised to the carrier controls. Two independent experiments, n=10 for carrier-treated mice, n=10 for sonidegib treated mice, unpaired Mann-Whitney test, mean ± SEM.
**(N)** Mean of number of CD8+ cells/ $mm^2$ in 100μm-wide zones from the tumour surface (-100 to 0 μm) to the tumour centre (-3300 to -3200 μm) as quantified by HALO software. Two independent experiments, n=9 for carrier treatment mice, n=9 for sonidegib-treated mice. Mean is shown in a single gradient heatmap.
